# A Compound Enhancing Lysosomal Function Reduces Tau Pathology, Microglial Reactivity and Rescues Working Memory in 3xTg AD Mice

**DOI:** 10.1101/2025.11.09.687389

**Authors:** Zachary Mayeri, Georgia Woods, Anand Rane, Alex Kifle, Manish Chamoli, Shikha Shukla, Chaska C. Walton, Julie K. Andersen

## Abstract

**Background:** Recent advancements in Alzheimer’s disease (AD) therapeutics have validated the use of amyloid beta (Aβ)-clearing antibodies, which reduce Aβ pathology but leave other disease hallmarks largely unaddressed. Since AD involves multiple pathological processes, additional strategies are needed to target complementary mechanisms. One such target is autophagy, a lysosomal mediated degradation pathway essential for cellular homeostasis that removes toxic protein aggregates and damaged organelles. This process is implicated in AD, as impaired lysosomal function promotes Aβ and tau accumulation. Our laboratory recently identified a novel natural mitophagy-inducing compound (MIC) that may serve as a therapeutic intervention for AD.

**Methods:** We evaluated the effects of MIC in aged 3xTgAD mice, a transgenic model displaying both Aβ and tau pathology. Mice received either standard diet or diet containing MIC beginning at age 4 months until 20 months on alternating weeks. Age-matched non-transgenic (NonTg) controls were included under standard and MIC-supplemented diets to assess compound effects during normal aging. Neuropathological changes were assessed using immunohistochemistry (IHC) for Aβ, phosphorylated tau (pTau), and microglial reactivity. Cognitive performance was evaluated using the Morris Water Maze (MWM) to assess spatial learning and memory and the Y-maze to measure working memory.

**Results:** At 20 months of age, our neuropathological assessment showed that 3xTgAD mice fed an MIC-supplemented diet had a significant reduction in pTau accumulation and microglial reactivity, although Aβ burden remained unchanged. At 15 months, MIC diet also improved spatial learning and memory in aged NonTg controls but not in 3xTgAD mice. However, in younger 8-month-old 3xTgAD mice, MIC restored working memory performance to NonTg levels, indicating an age-dependent therapeutic response.

**Conclusion:** MIC emerges as a potential modulator of tau pathology and neuroinflammation. As a naturally derived compound, MIC offers potential for combination therapy with FDA-approved Aβ-clearing antibodies, enabling a multimodal approach to AD treatment that addresses amyloid, tau, and microglia-related pathology.

## Introduction

Alzheimer’s disease (AD) and related dementias (ADRDs) currently affect an estimated 47 million people worldwide, a number projected to rise to 75 million by 2030 and 131 million by 2050, underscoring the urgent need for effective interventions [1]. AD is characterized by progressive neuronal degeneration, extracellular Aβ plaques and intracellular neurofibrillary tangles (NFTs) composed of hyperphosphorylated tau protein aggregates [2, 3]. While it is well documented that these protein aggregates and neuronal loss are indicators of disease progression, the upstream molecular mechanisms that trigger AD pathogenesis are not fully understood.

The amyloid hypothesis proposes that aggregation of Aβ initiates a cascade of neurotoxic events that culminate in synaptic dysfunction, neuroinflammation, and neuronal loss [4–8]. Aβ is generated through sequential proteolytic cleavage of amyloid precursor protein (APP), first by β-secretase and then by γ-secretase, whose catalytic subunit is encoded by the PSEN1 and PSEN2 genes [4]. Cleavage within the APP transmembrane domain releases Aβ peptides that aggregate into fibrils and eventually form extracellular amyloid plaques, a key pathological hallmark of AD [7, 9].

Recently, the U.S. Food and Drug Administration (FDA) approved several monoclonal antibodies targeting Aβ, designed to facilitate microglial clearance of amyloid plaques [10–13]. Although these therapies effectively reduce amyloid burden, their impact on cognitive decline remains modest, underscoring the need for complementary or alternative approaches that address additional mechanisms underlying Alzheimer’s pathology [14–16].

Alongside Aβ pathology, tau pathology represents a defining hallmark of AD [2, 3]. In AD, Tau proteins become hyperphosphorylated and disassociate from microtubules causing microtubules to disassemble [17]. Axonal pTau aggregates can reach the soma and form paired helical filaments (PHFs) leading to the development of NFTs [18]. It is well documented that the somatodendritic re-localization of pTau and subsequent aggregation into NFTs is what contributes to neuronal dysfunction and cell death associated with AD [19, 20].

Tau pathology is also targeted in AD clinical trials [21]. Tau targeting compounds in phase I clinical trials include APNmAb005, MK02214, and NIO752, in phase II BII080, Bepranemab, E2814, and JNJ-63733657, and in phase 3 E2814 [21, 22]. Tau-targeting antibodies are thought to act by preventing the trans-synaptic spread of toxic tau species [22]. ASO (e.g. BII080) acts by inhibiting tau RNA into tau proteins in neurons, thereby reducing total tau [21, 23] However, there remain no FDA approved interventions that have been shown to enhance intracellular protein clearance of pTau aggregates.

Cognitive dysfunction in AD extends beyond canonical Aβ and tau pathologies to encompass pathological microglial activation, characterized by a pronounced shift from a resting surveillant phenotype to a pro-inflammatory reactive state [24, 25] This phenotypic transition has been mechanistically linked to chronic exposure and aberrant sensing of AD associated pathogenic proteins.

Microglia mediated neuroinflammation in AD has also been targeted for therapeutic intervention [26, 27]. OLT1177 an oral NLRP3 inflammasome inhibitor, acts upstream to prevent IL-1β maturation from microglia [26]. XPro1595 (Pegipanermin), a TNF-neutralizing protein in Phase 2 trials, demonstrated decreased neuroinflammation biomarkers and improved white matter structure in mild-to-moderate AD patients [27]. These complementary therapeutic approaches highlight the emerging recognition that modulating pathological microglial immune activation represents a promising strategy for slowing cognitive decline in Alzheimer’s disease.

Cellular protein and organelle clearance are mediated by the autophagy-lysosomal pathway (ALP), which relies on fusion of autophagosomes with lysosomes [28–30]. Autophagic and lysosomal dysfunction are well documented in AD studies [31–36]. Transcription factor EB (TFEB) is established as the master regulator of the ALP [37]. In pursuit of therapeutic strategies to modulate TFEB activity, we conducted a natural product screen in N27 rat dopaminergic neural cells and identified a compound we called mitophagy inducing compound (MIC) via its ability to enhance mitochondrial turnover via increased TFEB promoter activity 6-fold. Validation in *C. elegans* demonstrated that MIC increases activity of HLH-30—the TFEB ortholog in worms— enhances lysosomal function, reduces proteotoxicity and extends lifespan as well as lysosomal function in mammalian cells [38]. Here we report that in aged 3xTg AD mice, longitudinal dietary supplementation with MIC resulted in significant reductions in pTau, nuclear localized pTau, and perinuclear pTau, without affecting Aβ pathology.

Furthermore, MIC administration led to significant reduction in microglia reactivity in aged 3xTg AD mice while having no effect on microglia reactivity in aged nonTg mice. MIC was found to improve working memory independently of locomotive behavior at the 8-month timepoint in the 3xTg AD mice, which was lost at later disease stages. This corresponded with a lack of effects on spatial memory performance in 3xTg AD mice. However, MIC did elicit significant improvement in spatial memory in aged nonTg mice.

## Materials and Methods

### Animals

All experiments involving mice were approved by the Buck Institute for Research on Aging Institutional Animal Care and Use Committee (IACUC) and were conducted in compliance with all relevant ethical regulations for animal research. 3xTg-AD female mice and B6129SF2/J non-transgenic female mice were obtained from The Jackson Laboratory (Ellsworth, ME). All animals were housed in the Buck Institute vivarium (Novato, CA) under standard conditions for social housing and environmental enrichment, in accordance with institutional animal care guidelines.

### MIC Treatments

From birth to 4 months of age, all mice were fed a vivarium diet (irradiated food rodent diet, Cat TD-2918, Envigo Teklad, Madison, WI). Starting at 4 months, 3x-Tg AD mice (n=15) and non-Tg mice (n=15) were fed either a control (CTRL) diet (D110700GI, AIN-93G containing sterile casein; Dyets, Inc, Bethlehem, PA) or MIC (25 mg/kg MIC in AIN-93G containing casein, 104252GI on alternating weeks with fed of CTRL diet until euthanization at 20 months (Fig. 1a).

**Figure 1:**
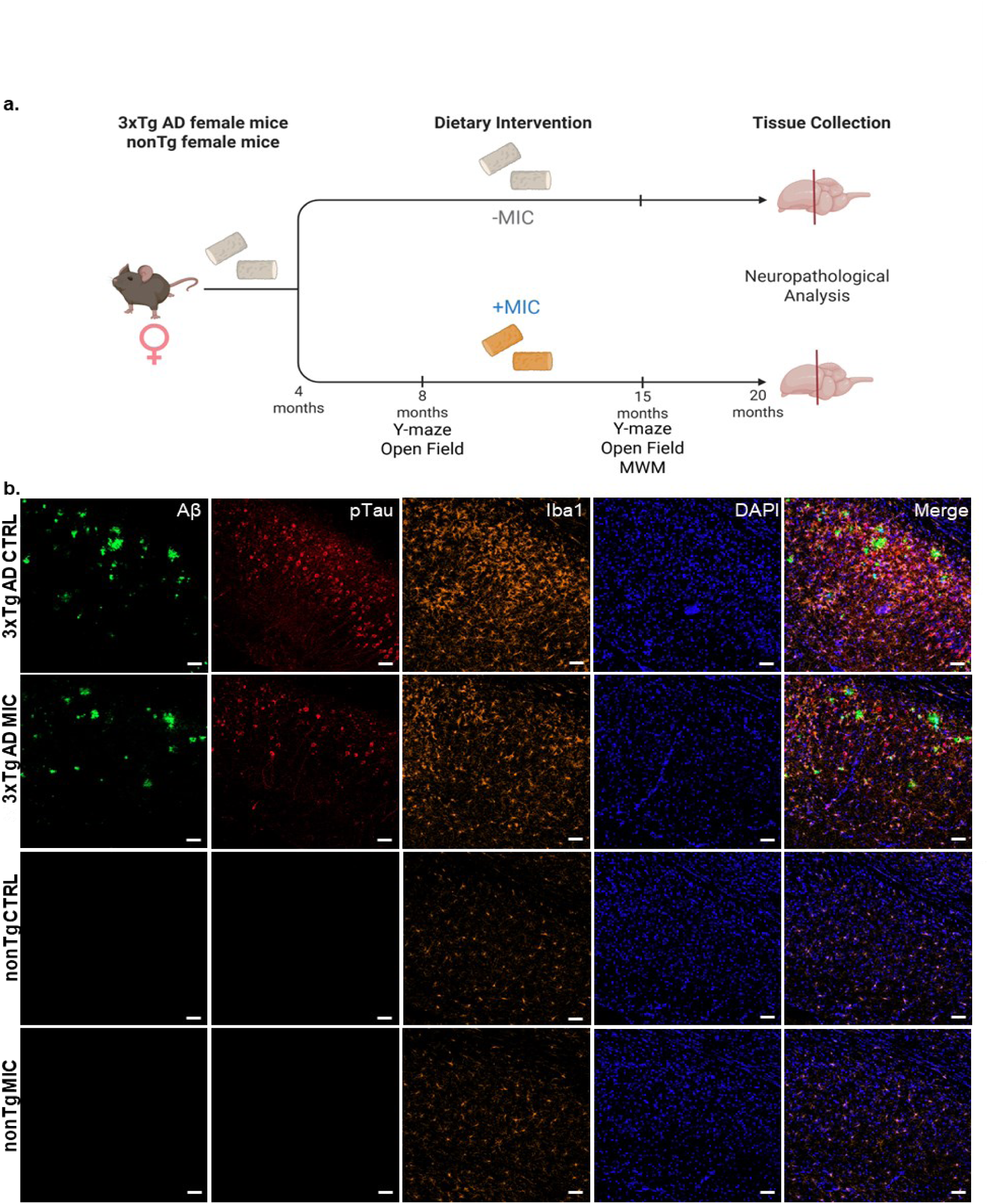
Project timeline and neuropathological analysis of aged 3xTg AD mice and nonTg mice. **(a)** Schematic depicting female 3xTg AD mice and female nonTg mice started on a normal chow vivarium diet from birth to 4 months. At 4 months, mice were switched to a CTRL diet or MIC diet every other week with CTRL diet on off weeks until 20 months when all mice were euthanized for neuropathological analysis. Longitudinal behavioral analyses were performed as described at 8 and 15 months of age. **(b)** Immunofluorescence staining of regionally matched 40 μm thick slices of the dorsal subiculum from the caudal hippocampus of 20 month-old 3xTg AD and nonTg mice fed on CTRL and MIC diets. Visualization of Aβ, (6e10, [1–16]) (green), pTau (AT8 [Serine 202/Threonine 205]) (red), microglia [Iba1] (orange) and cell nuclei [DAPI] (blue). Scale bar = 50 μm.

### Y-maze

Working memory was assessed in the Y-maze test following established protocols [39, 40]. The Y-maze apparatus consisted of with three symmetrical arms measuring 40 × 8 × 15 cm (length × width × height). Mice were acclimated to the testing room for 1 hour prior to testing. Each mouse was positioned in the center zone and allowed unrestricted exploration for 10 minutes. Arm entries and alternations were automatically tracked and quantified using the EthoVision XT software (Noldus Information Technology, Leesburg, VA; RRID: SCR_000441). In accordance with established protocols [39], mice that did not move throughout the duration of the test were re-trialed. Mice that remained still during trial and retrial were excluded from the spontaneous alternation analysis to assess working memory, but still included in the arm entry analysis to assess locomotion.

### Open Field

The Open Field Test was performed as previously described by our laboratory with small variations [41]. Locomotory and exploratory behavior was assessed using the TruScan photobeam apparatus (Coulbourn Instruments, Holliston, MA, USA). Mice were placed in the open field chamber and allowed to explore freely for 15 minutes. The distance traveled during the 15-minute period was quantified using TruScan 2.0 software (Coulbourn Instruments).

### Morris Water Maze (MWM)

The MWM was adapted from previously described procedures [42, 43]. The MWM consisted of a circular pool (122 cm diameter, 50 cm height) filled with water maintained at 22 ± 1 °C and rendered opaque with non-toxic white paint. A hidden escape platform was submerged just below the surface in the center of the northwest quadrant. The testing room contained fixed distal visual cues that remained unchanged throughout the experiment. Mice underwent acquisition training with two trials per day for five consecutive days. On day 1, trial 1, mice were placed at the southernmost point of the pool and given 60 seconds to locate the hidden platform. If unsuccessful, they were guided to the platform and allowed to rest for 10 seconds. Trial 2 followed the same procedure. On days 2–5, trial 1 also began from the southernmost point, while trial 2 began from the easternmost point, following the same timing and placement protocol as day 1. Twenty-four hours after the final training session, the platform was removed, and mice underwent one 30-second probe trial in the morning followed by a rest and then a 60-second probe trial in the afternoon, both initiated from the southeasternmost point. The number of crossings over the platform’s previous location was recorded. Behavior was recorded and tracked using EthoVision XT software (Noldus Information Technology, Leesburg, VA; RRID: SCR_000441).

### Tissue Collection and Histology Preparation

Mice from all 4 conditions were euthanized in a CO_2_ chamber at 20 months of age. Mice were exsanguinated via transcardial perfusion with 1X phosphate buffered saline (PBS). These timepoints were selected to enable a standardized comparison for neuropathological analysis as previously described [44]. Brains were dissected and stored at 4° C in 4% paraformaldehyde for 48 hours and then transferred to PBS. Tissue Sectioning

For tissue sectioning, the brains were embedded in 4% agarose at room temperature or in Optimal Cutting Temperature (OCT) compound (4583, Tissue-Tek, Torrance, CA) and frozen at −80 ° C. They were sectioned at a thickness of 40 μm per section along the coronal plane, from rostral to caudal using a cryostat (CM 1950, Leica) respectively. Sections were collected in ice cold tris buffer saline (TBS); and transferred to a 48-well plate containing cryobuffer (0.1 M Potassium Acetate, 40% Ethylene Glycol, 1% Poly Vinyl Pyrrolidone in PBS, Sigma-Aldrich, Darmstadt, Germany). Sections were stored at −20° C.

### Immunohistochemistry

Sections from each mouse were regionally matched, hemisected and selected for staining. All sections were stained with the following protocol at room temperature unless otherwise noted. Free-floating sections were washed three times with TBS to remove the cryobuffer. For antigen retrieval, individual sections were placed in 2 mL Eppendorf tubes containing 1 mL of citrate buffer (6.0 pH; sodium citrate tribase dehydrate, citric acid anhydrous, Sigma-Aldrich), tightly wrapped with parafilm and then put into a 90° C water bath for 10 minutes. Sections were then washed three times in TBS, permeabilized for 1 h in 0.2% Triton X-100 (Thermo Fisher Scientific, Waltham, MA) in TBS, and blocked for 1 h in 5% normal donkey serum (NDS; 017-000-121, Jackson ImmunoResearch, West Grove, PA) prepared in 0.2% Triton-TBS. Tissue sections were incubated overnight at 4 °C with primary antibodies diluted in 5% NDS and 0.2% Triton-TBS. The following day, sections were washed three times with TBS containing 0.1% Tween-20 (TBS-Tween, Sigma-Aldrich) then incubated with secondary antibodies diluted in TBS-Tween for 1 h at room temperature. Sections were subsequently washed in TBS-Tween for 2 h to remove unbound antibody. For conjugated antibody staining, sections were incubated in TBS-Tween for 1 h at room temperature following completion of the primary and secondary antibody steps. After final washes in TBS-Tween, sections were counterstained with DAPI (1:10,000; Invitrogen) for 10 min, mounted onto microscope slides (VWR Premium Superfrost Plus Microscope Slides, Avantor, Visalia, CA) and coverslipped (VWR Rectangular Micro Cover Glasses, Avantor) using ProLong Gold Antifade Mountant (P36930, Invitrogen).

### Antibodies

Primary antibodies: anti-phospho-Tau Ser202, Thr205 AT8 mouse antibody (1:200 dilution) (MN1020, Invitrogen), anti-Iba1 (1:200 dilution) (019-19741, FUJIFILM Wako Diagnostics, Mountain View, CA, US), and Alexa Fluor 488 conjugated anti-β-amyloid 1-16 6E10 (1:200) (Biolegend, 803019). Secondary antibodies: donkey anti-mouse Alexa-Fluor 647 (A31471, Invitrogen) and donkey anti-rabbit Alexa-Fluor 555 (A31572, Invitrogen).

### Confocal Imaging and Image Processing

Images were acquired with an LSM-980 confocal microscope (Zeiss, Jena, Germany.) with a 20x objective lens using Zen Image Acquisition software (Zeiss, RRID: SCR_013672). Z-stack tile scan images of 707 um^2^ of the dorsal subiculum (Bregma - 3.16mm, Supplemental Fig. 1) directly adjacent to but not in view of the dorsal CA1 pyramidal cell layer were acquired. IMARIS (Zurich, Switzerland, RRID: SCR_007370) was used for 3D visualization, video generation, and processing of 3D confocal images. A standardized volume was selected from within the tile scans to ensure equivalent regions were compared in the z-plane. Standardized 3D regions of interest (ROI) for Aβ, pTau, Iba1, and DAPI-positive voxels (volumetric pixel) were then generated using the surface function of IMARIS. The algorithms used to generate the ROIs were standardized for each replicate to eliminate user bias. **Aβ and pTau volume analysis:** The total volume of Aβ and pTau was obtained from the sum volume of 3D ROIs calculated from voxels positive for each marker. **Aβ and Iba1 protein quantification:** Total protein levels of Aβ and Iba1 were quantified in arbitrary units to assess amyloid burden and microglial reactivity, respectively. For each sample, the total sum voxel fluorescence intensity within the 3D ROIs corresponding to Aβ or Iba1 immunoreactivity was summed to obtain total protein levels. Additionally, mean microglial protein levels were determined by averaging the mean voxel fluorescence intensity within individual Iba1-positive 3D ROIs. **Percent pTau positive cells:** 3D ROI based on DAPI were used to identify nuclei and 3D ROI based on the pTau channel were used to identify pTau-positive structures. The distance function in IMARIS was used to assess distances between the ROI types. DAPI ROI in direct contact (distance = 0 μm) to pTau ROI was used to identify pTau positive nuclei. The percentage of pTau-positive nuclei was used as a proxy for pTau-positive cells. **Nuclear and perinuclear localized pTau:** pTau positive nuclei generated as described above were used to partition perinuclear and nuclear pTau 3D ROI. Briefly, pTau signal within DAPI-positive neuclei was categorized as nuclear pTau and the remaining pTau that was < 2 μm from the edge of nuclei was categorized as perinuclear pTau. The sum voxel pTau fluorescence intensity for nuclear or perinucluear pTau ROIs was normalized to the total number of nuclei assessed per image and used as proxies of total nuclear and perinuclear pTau protein.

### Statistical Analysis

Statistical analyses were conducted using IBM SPSS Statistics (Chicago, IL, RRID: SCR_002865) and data was graphed using GraphPad Prism (San Diego, CA, RRID: SCR_002798). Independent samples t-tests and two-Way ANOVAs with post hoc Tukey’s multiple comparisons were run to compare groups. Statistical significance was set at p < 0.05, two-tailed. Assumptions tests were carried out to detect extreme outliers (box plots), normality (Shapiro-Wilk test, p > 0.05), and homogeneity of variances (Levene’s test, P > 0.05). When statistical assumptions were violated, appropriate data transformations were applied, and all assumption tests were subsequently re-evaluated to confirm compliance. Non-parametric Kruskal–Wallis tests with Bonferroni-corrected all-pairwise comparisons on ranks were conducted when data transformations failed to meet the assumptions required for parametric analyses. For behavioral data, figures display non-transformed values to facilitate visual interpretation. Comprehensive results for both transformed and non-transformed datasets—including assumption tests, omnibus analyses, and post hoc comparisons—are provided in Supplemental Tables 1– 6. Exact p-values are reported for all outcomes, regardless of significance.

## Results

### Neuropathology in 3xTg AD and nonTg Mice

3xTg AD mice are a widely used model of AD that develop age-dependent pathology, originally generated by Oddo et al. through co-injection of Presenilin 1 Methionine-146-Valine (PS1M146V) and Amyloid Precurser Protein Swedish (APPSwe) transgenes, which are linked to familial Alzheimers Disease (fAD), as well as the tauP30IL transgene, which is linked to frontotemporal dementia (FTD) [45, 46]. In these mice, Aβ plaques progressively accumulate in the hippocampus (80% of mice at 6 months, 100% of mice at 12 months) and are found in the subiculum, CA1 field, and entorhinal cortex, while pTau localizes both intraneuronally and extracellularly in pyramidal cells, increasing in an age-dependent manner starting as early as 4 months of age [44, 47]. Only female mice were used because they develop Aβ pathology [48] and pTau pathology [44] earlier and in greater amounts than their male siblings. We used age-matched female B6129SF2/J nonTg groups as controls.

MIC or control diet begun at 4 months for the 3xTg AD (CTRL n = 15, MIC n = 15) and nonTg control (CTRL n = 15, MIC = 15) groups to determine the drug’s ability to suppress the age-dependent accumulation Aβ ant tau pathology as well as microglia reactivity (Fig. 1a). Five mice from 3xTg AD CTRL diet group, 4 mice from the nonTg CTRL diet group, 8 mice from the 3xTg AD MIC diet group, and 2 mice from the nonTg MIC diet died from natural causes and were not included in the studies of neuropathology.

The remaining mice were humanely euthanized at 20 months of age and immunohistochemistry (IHC) was performed on 40 μm-thick regionally matched hippocampal slices from the dorsal subiculum (Sup. Fig. 1), critical for spatial memory and learning in mice [49]. Tissue sections were immunolabeled to detect Aβ using the 6E10 antibody, microglial activation using Iba1 antibodies, and hyperphosphorylated tau (Ser202/Thr205) using the AT8 antibody (Fig. 1b) [50–52]. Cell nuclei were counterstained with DAPI (Fig. 1b). As expected, the comparison between 3xTgAD and nonTg mice confocal microscope images revealed substantial Aβ plaque deposition (6E10), pTau accumulation (AT8), and microglia reactivity (Iba1) in 3xTg AD mice regardless of the diet (Fig. 1b).

### MIC does not reduce Aβ neuropathology in 3xTg AD mice

In 20-month-old 3xTgAD mice, confocal imaging revealed readily identifiable large, compact plaques with strong Aβ signal (red boxes) as well as more diffuse aggregates exhibiting fainter Aβ labeling (yellow boxes) in both MIC- and control (CTRL)-fed groups (Fig. 2a; Sup. Fig. 2).

**Figure 2:**
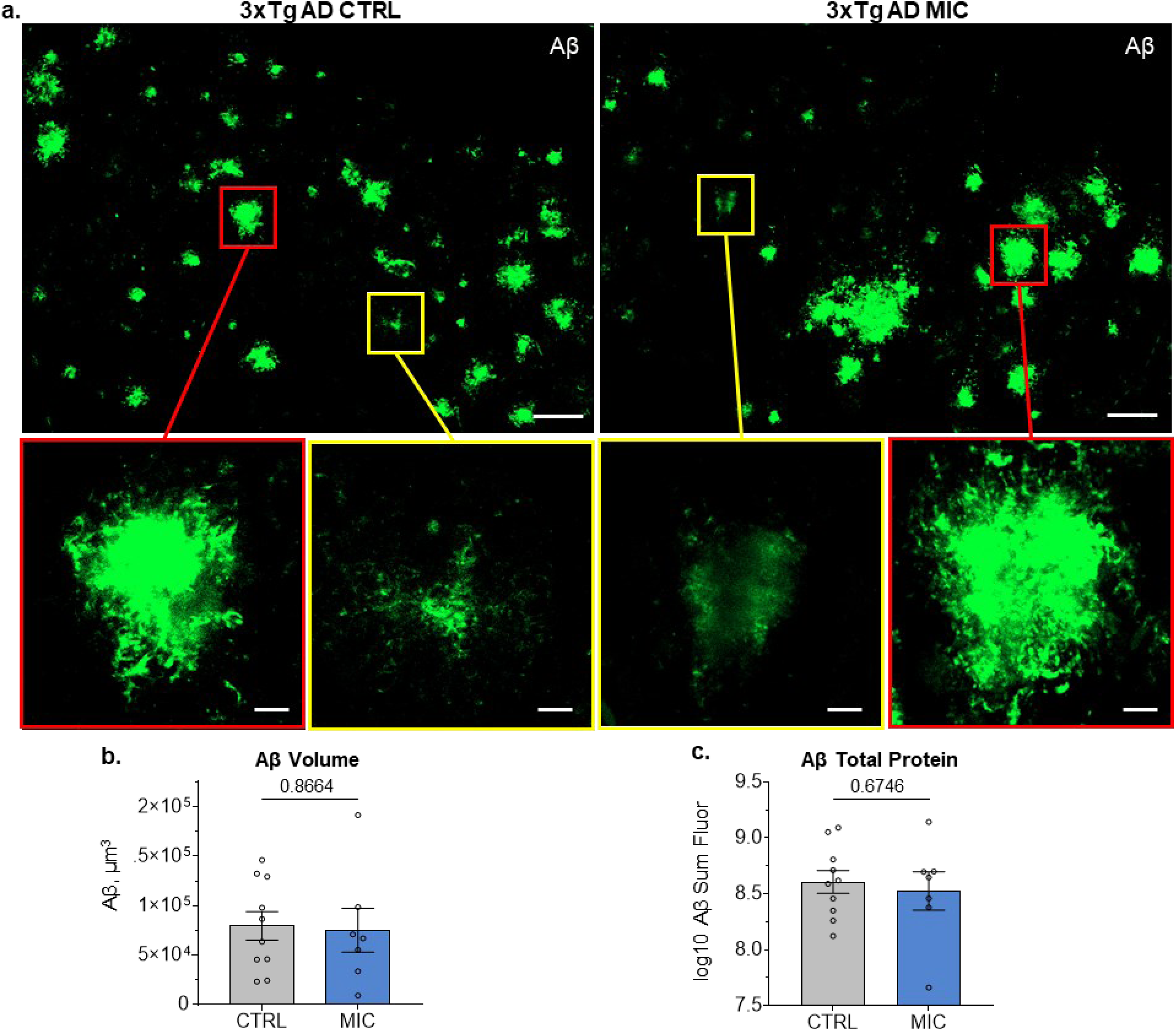
Aβ pathology in dorsal subiculum of 3xTg AD mice is not altered with MIC treatment. **(a)** Representative images from 3xTg AD mice on CTRL and MIC diet Aβ plaques in the dorsal subiculum of 3xTg AD mice. Scalebar, 50 μm. Boxes in top image are expanded bellow, showing details of compact (red boxes) and diffuse (yellow boxes) Aβ aggregates. Scalebar, 10 μm. Independent *t*-Test for **(b)** Aβ volume (μm^3^) and **(c)** log10 transformed sum fluorescence intensity. Log10 transform was performed to meet the assumption of normality. For full statistical analysis see Sup. Table 1. CTRL (n = 10), MIC (n = 7). For Bars display mean ± SEM.

To evaluate potential differences in the overall abundance of Aβ aggregates, 3D ROIs were generated using Imaris image analysis software to quantify the total Aβ-positive volume within dorsal subiculum sections. This volumetric analysis measures aggregate abundance independent of fluorescence intensity, allowing direct comparison of Aβ burden between MIC-and CTRL-fed mice regardless of variations in signal brightness. For the comparison between 3xTgAD CTRL and MIC diet groups, an independent *t*-Test revealed no significant difference in total Aβ volume (t(15) = 0.171, p = 0.8664; Fig. 2b; Sup. Table 1a), indicating that MIC did not reduce overall Aβ aggregate load.

To further assess potential changes in Aβ protein abundance, the total Aβ fluorescence signal—a relative measure of total Aβ content—was quantified within the same ROIs. Consistent with the volumetric findings, no significant difference was observed between 3xTgAD CTRL and MIC diet groups (t(15) = 0.428 p = 0.6746; Fig. 2c; Sup. Table 1b). Taken together, MIC did not reduce the total volume of Aβ aggregates nor the abundance of Aβ within aggregates over the 16-month treatment period.

### MIC significantly reduces pTau in 3xTg AD mice

In contrast to Aβ, confocal imaging revealed visible differences in pTau labeling between 3xTgAD CTRL and MIC diet groups, with noticeably reduced pTau signal in MIC-treated mice (Fig. 3a; Sup. Fig. 3). To further quantify this observation, we performed the same volumetric analysis previously applied to Aβ aggregates. The total pTau-positive volume was significantly reduced in MIC-treated mice compared to CTRL animals (t(15) = 3.333, p = 0.0045; Fig. 3b; Sup. Table 2a), indicating a reduction in pTau-positive structures following MIC supplementation.

**Figure 3:**
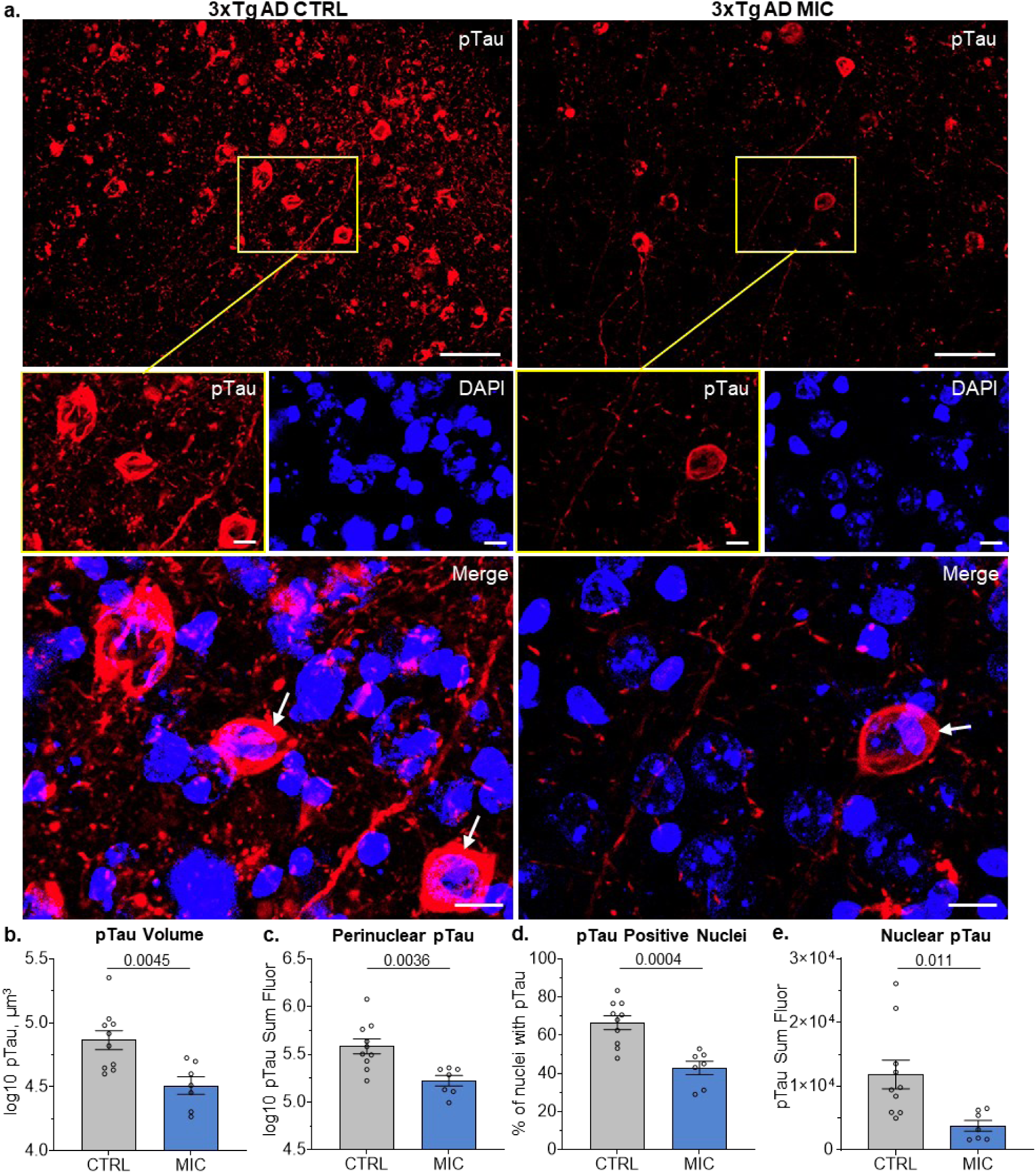
Tauopathy in the dorsal subiculum of 3xTg AD mice is reduced with MIC treatment. **(a)** Representative images (top) from 3xTg AD mice on the CTRL and MIC diet visualizing pTau (AT8) in the dorsal subiculum. Scale bar, 50 μm. Detailed images (middle) of square regions in top images showing pTau and DAPI as separate channels as well as their merged image (bottom). Scale bar, 10 μm. **(b)** Independent *t*-Tests for pTau volume, **(c)** perinuclear pTau fluorescence, **(d)** percent-positive pTau nuclei, and **(e)** nuclear pTau fluorescence. **(b, c, e)** Log10 transforms were performed to meet ANOVA assumptions. For full statistical analysis see Sup. Table 2. CTRL (n = 10), MIC (n = 7). Significance, p < 0.05. Bars display mean ± SEM.

To determine whether this volumetric decrease was accompanied by a reduction in intracellular pTau protein levels in live cells, we next analyzed fluorescence signal intensity within cellular compartments. Specifically, we identified pTau signal within 2 μm of DAPI-positive nuclei (Fig. 2a, white arrows), representing pTau localized to live cells. The signal was further subdivided into perinuclear and nuclear pTau, depending on whether it overlapped with DAPI staining or not (Sup. Video 1).

Perinuclear AT8+ pTau represents early detectable somatic accumulation of pTau, marking a critical transition point where tau escapes its normal axonal compartment and begins aggregating in the neuronal soma before progressing to mature NFTs [53, 54]. Perinuclear fluorescence intensity was highly significantly decreased in MIC-treated mice (t(15) = 3.451, p = 0.0036) (Fig 3c; Sup. Table 3b), evidencing MIC was reducing signal associated to pathological tau in live cells.

Under normal conditions, nuclear tau regulates DNA stability and heterochromatin structure in both human iPSC derived neurons and mouse primary cortical neurons [55–57]. In our 3xTg AD mice, we observed the colocalization of AT8+ pTau with DAPI, indicating pTau is localizing inside of cell nuclei. For nuclear pTau, we quantified both the percentage of cells positive for nuclear pTau as well as the intensity of nuclear pTau. Statistical analysis revealed that MIC treatment highly significantly reduced both percentage of pTau-positive nuclei (t(15) = 4.487, p = 0.0004) (Fig 3d, Sup. Table 3c) and total nuclear pTau fluorescence intensity (t(15) = 2.882, p = 0.0011) (Fig 3e; Sup. Table 3d).

These results demonstrate that chronic MIC administration significantly attenuated tau pathology in 3xTgAD mice. Notably, the reduction encompassed both perinuclear and nuclear pTau compartments, indicating that MIC mitigates intracellular tau accumulation and may act, at least in part, by reducing tau burden within viable neurons.

### MIC significantly reduces microglia reactivity in 3xTg AD mice

In contrast with Aβ and pTau absent from nonTg, Iba1 signal was readily detected in both 3xTg AD and nonTg mice (Fig. 1b). Iba1 is calcium-binding protein that serves as a widely used marker for detecting microglia throughout the brain in health and pathology. Iba1 is upregulated during microglial activation and plays crucial roles in membrane ruffling and cytoskeleton reorganization [51, 58]. Accordingly, Iba1 staining was analyzed in both nonTg and 3xTgAD mice to assess microglial distribution and activation status.

Confocal images of nonTg and 3xTg AD mice revealed distinct morphological differences between experimental groups (Fig. 4a, Sup. Fig. 4a, b, Supp Fig. 5). In nonTg mice, microglia display the characteristic ramified morphology found in healthy aging brains, with thin, highly branched processes extending radially from small cell bodies (Sup. Fig. 5) [59, 60]. Compared to nonTg mice, 3xTg AD mice show markedly different microglial morphology with increased Iba1 intensity and thicker processes, characteristic of the activated state observed in the AD mouse models with amyloid pathology [44, 47, 61, 62] (Fig. 4a, Sup. Fig. 4a & 4b).

**Figure 4:**
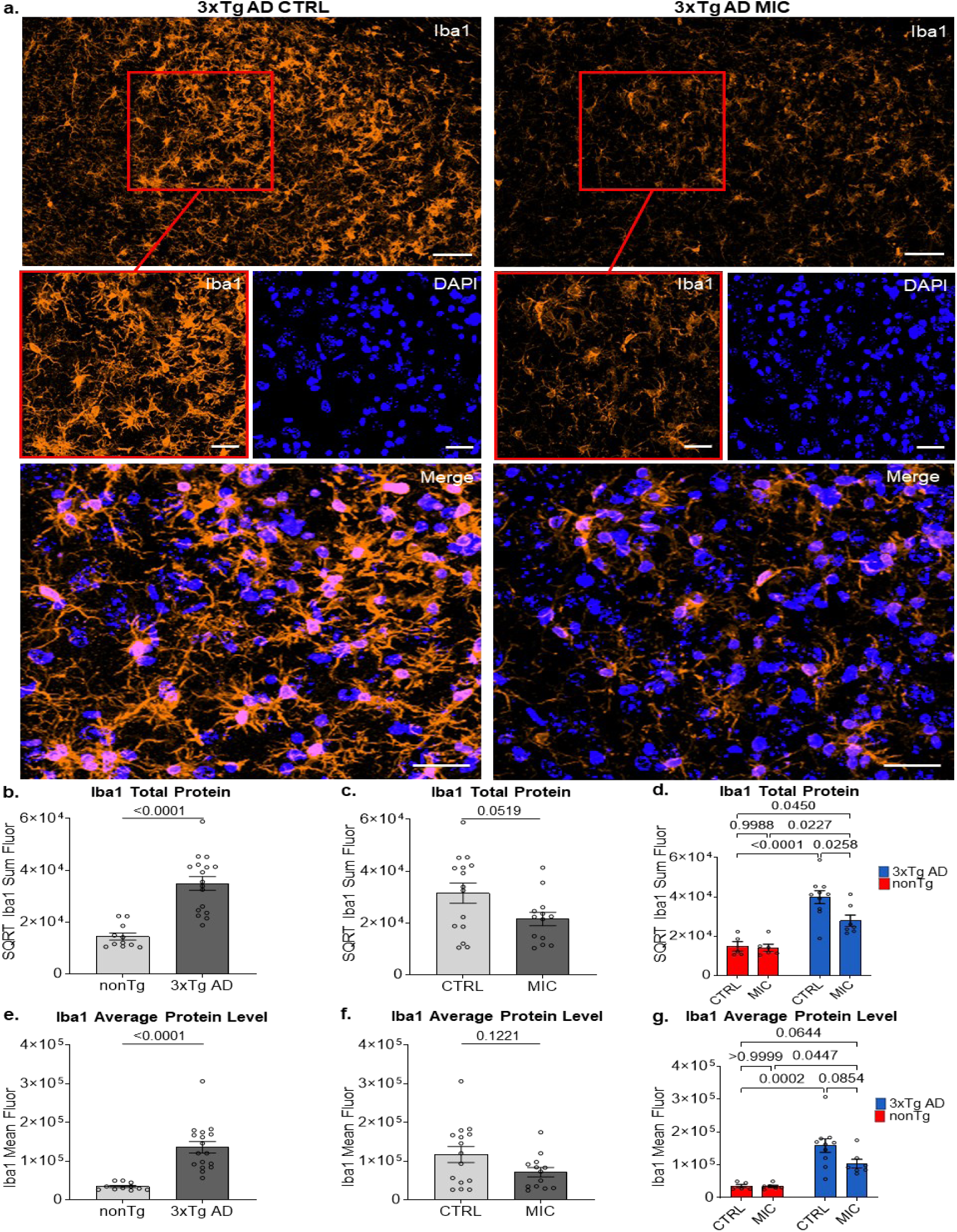
MIC treatment reduces microglia reactivity in the dorsal subiculum of 3xTg AD mice. **(a)** Representative images (top) from 3xTg AD mice on the CTRL and MIC diets visualizing Iba1 in the dorsal subiculum. Scale bar, 50 μm. Detailed images (middle) of square regions in top images showing Iba1 and DAPI as separate channels as well as their merged image (bottom). Scale bar, 20 μm. **(b)** Genotype main effects (nonTg & 3xTg AD), **(c)** diet (CTRL & MIC) ANOVA main effects, and **(d)** Tukey multiple comparisons for sum Iba1. Iba 1 fluorescent intensity was square root transformed to meet ANOVA assumptions. **(b)** Genotype main effects (nonTg & 3xTg AD), **(c)** diet (CTRL & MIC) ANOVA main effects, and **(d)** Tukey multiple comparisons for mean Iba1 fluorescent intensity. For full statistical analysis see Sup. Table 3. nonTg CTRL (n = 5), nonTg MIC (n = 6), 3xTg AD CTRL (n = 10), 3xTg AD MIC (n = 17). Significance, p < 0.05. Bars display mean ± SEM.

**Figure 5:**
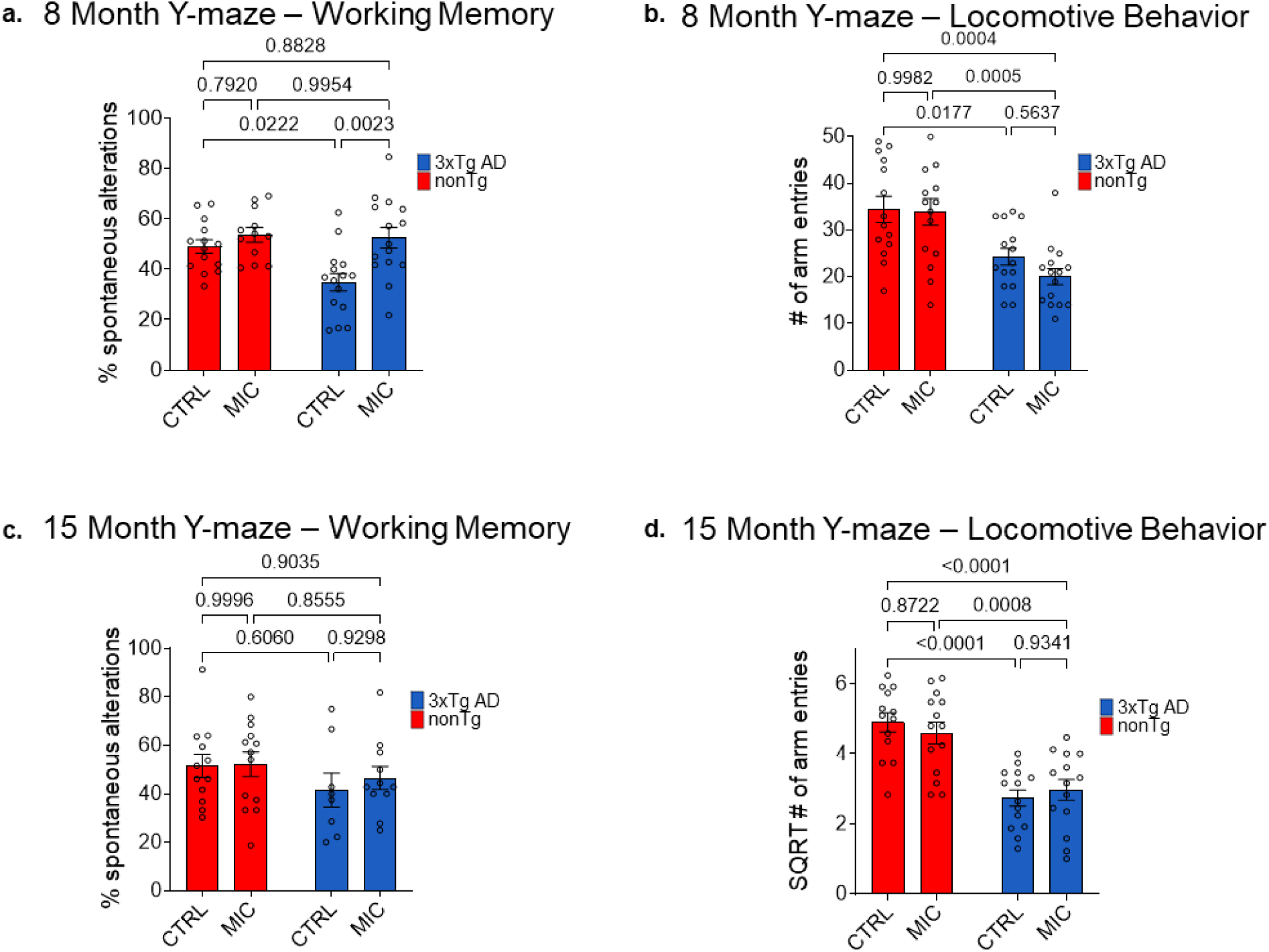
Y-maze performance in 8-month and 15-month old female mice. (**a, b**) Percent spontaneous alternation (**a**) and number (#) of total arm entries (**b**) at 8 months of age. Animals that did not perform any arm entries were excluded from the percent alternation analysis. (**a**) nonTg CTRL (n = 14), nonTg MIC (n = 12), 3xTg AD CTRL (n = 15), 3xTg AD MIC (n = 15). (**b**) For total arm entries, all animals were included. nonTg CTRL (n = 14), nonTg MIC (n = 15), 3xTg AD CTRL (n = 15), 3xTg AD MIC (n = 15). (**c, d**) Percent spontaneous alternation (**c**) and number (#) of total arm entries (**d**) at 15-months of age. Animals that did not perform any arm entries were excluded from the percent alternation analysis. (**c**) nonTg CTRL (n = 12), nonTg MIC (n = 13), 3xTg AD CTRL (n = 8), 3xTg AD MIC (n = 11). (**d**) For total arm entries, all animals were included. nonTg CTRL (n = 13), nonTg MIC (n = 14), 3xTg AD CTRL (n = 14), 3xTg AD MIC (n = 14). (**a-d**) Tukey multiple comparisons. For full statistical analysis see Supplemental Table 5. Significance, *p* < 0.05. Bars display mean ± SEM.

For quantitative comparison of Iba1 across genotypes and diet groups, we first measured the summed fluorescence intensity of Iba1 as a relative indicator of total microglial reactivity. A two-way ANOVA revealed no significant interaction between genotype and diet (*F*_(1, 24)_ = 3.300, p = 0.082, partial η^2^ = 0.121; Sup. Table 3a). An interaction assessment analyzes whether the effect of one independent variable on the outcome depends on the level of another independent variable. As expected, there was a highly significant main effect of genotype, with Iba1 levels markedly higher in 3xTgAD mice than in nonTg controls (*F*_(1, 24)_ = 39.152, p < .001, partial η^2^ = 0.620; Fig. 4b, Sup. Table 3a). Main effects represent the independent influence of each factor on the outcome variable, averaged across all levels of other factors. A non-significant trend was also observed for diet, suggesting a possible reduction in Iba1 signal in MIC-fed mice compared to CTRL-fed mice (*F*_(1, 24)_ = 4.185, p = 0.052, partial η^2^ = 0.148; Fig. 4c Sup. Table 3a). Multiple comparisons followed to identify potential biologically meaningful effects. Tukey’s multiple comparison test revealed a significant reduction in total Iba1 signal in MIC-fed relative to CTRL-fed 3xTgAD mice (p = 0.0258), an effect not observed in nonTg animals (p = 0.9988; Fig. 4d, Sup. 3a). These findings indicate that the trend toward reduced Iba1 levels in the main diet effect (p = 0.052) was primarily driven by the MIC-associated decrease in microglial reactivity within the 3xTgAD group (p = 0.0258), indicating a potentially biologically relevant effect of MIC specifically in 3xTgAD mice.

We next analyzed the mean fluorescence intensity of Iba1 per 3D ROI as a proxy of average microglial activation. As with the total Iba1 signal, the two-way ANOVA showed no significant interaction between genotype and diet (*F*_(1, 24)_ = 2.412, p = 0.133, partial η^2^ = 0.091; Sup. Table 3b) and a highly significant main effect of genotype (*F*_(1, 24)_ = 30.912, p < 0.001, partial η^2^ = 0.563; Fig. 3e; Sup. Table 3b). In contrast to the summed Iba1 signal, the mean Iba1 intensity did not display a trend for the main effect of diet (*F*_(1, 24)_ = 2.569, p = 0.122, partial η^2^ = 0.097, Fig. 4f, Sup. Table 3b). Follow-up Tukey’s multiple comparison tests revealed no significant differences between MIC- and CTRL-fed 3xTgAD mice (p = 0.0854) or between diet groups in nonTg mice (p > 0.9999; Fig. 4g, Sup. Fig. 3b).

In summary, MIC supplementation did not significantly affect either total or mean Iba1 signal in nonTg mice. In 3xTgAD mice, the MIC diet produced a potentially biologically meaningful reduction in total Iba1, but not in mean intensity. This pattern may suggest that MIC could reduce the number of reactive microglia, while maintaining the same average activation level in the microglial population.

### MIC rescues working memory in 8 but not 15 month-old 3xTg AD mice

While the neuropathological analyses described above were conducted in 20-month-old mice, behavioral testing using the Y-maze spontaneous alternation task was performed at both 8 and 15 months of age to enable a longitudinal assessment of MIC effects. The Y-maze spontaneous alternation test assesses spatial working memory in rodents based on their innate tendency to explore novel environments [63, 64]. Mice alternate between maze arms at rates significantly above chance with this behavior dependent on hippocampal integrity [65, 66]. Following established protocols, we measured both spontaneous alternation percentage (working memory) and arm entries (locomotor activity) to distinguish cognitive deficits from behavioral confounds [39, 40].

At 8 months of age, a two-way ANOVA revealed no significant interaction between genotype and diet on the percentage of spontaneous alternations (*F*_(1, 52)_ = 3.579, p = 0.064, partial η^2^ = 0.064, Sup. Table 4a). In contrast, the main effects for genotype (*F*_(1, 52)_ = 4.971, p = 0.030, partial η^2^ = 0.087) and diet (*F*_(1, 52)_ = 10.541, p = 0.002, partial η^2^ = 0.169, Sup. Table 4a) were both significant. Follow-up Tukey’s multiple comparisons indicated a significant increase in spontaneous alternation performance in MIC-fed 3xTgAD mice compared to CTRL-fed counterparts (p = .0023), rescuing performance to levels that were not significantly different from nonTg controls (p = 0.8828, Fig. 5a, Sup. Table 4a). There were no significant differences observed between diet groups in nonTg mice (p = 0.7920, Fig. 5a, Sup. Table 4a). Thus, despite the significant main effects for diet (p = 0.002), post-hoc analysis suggest that the effect of MIC was primarily driven by a rescue of working memory performance in 3xTgAD mice.

To control for potential confounding differences in locomotor activity that may influence the above reported Y-maze results, we analyzed the total number of arm entries, irrespective of alternation sequence. The two-way ANOVA revealed no significant interaction between genotype and diet (*F*_(1, 55)_ = 0.629, p = 0.431, partial η^2^ = 0.011) and no significant main effect of diet (*F*_(1, 55)_ = 1.074, p = 0.305, partial η^2^ = 0.019; Sup. Table 4b). In contrast, the main effect of genotype was highly significant (*F*_(1, 55)_ = 26.548, p < 0.001, partial η^2^ = 0.326; Sup. Table 4b), reflecting a reduction in total arm entries in 3xTgAD mice compared to nonTg controls. Post hoc multiple comparisons revealed no significant differences between MIC- and CTRL-fed mice within either genotype (3xTgAD: p = 0.5637; nonTg: p = 0.9982; Fig. 5b; Sup. Table 4b).

Overall, these results indicate that while 3xTgAD mice exhibited reduced locomotor activity relative to nonTg animals, MIC treatment did not alter movement behavior within either group. Therefore, the enhanced spontaneous alternation performance observed in MIC-fed 3xTgAD mice reflects a true improvement in working memory, rather than differences in general motor activity.

The Y-maze test was repeated at 15 months of age to assess whether the effects of MIC persisted at later stages. At this age, the two-way ANOVA revealed no significant interaction between genotype and diet (F_(1, 40)_ = 0.152, p = 0.698, partial η² = 0.004; Sup. Table 4c) and no significant main effects of either diet (F_(1, 40)_ = 0.278, p = 0.601, partial η² = 0.007; Sup. Table 4c) or genotype (F_(1, 40)_ = 2.116, p = 0.154, partial η² = 0.050; Sup. Table 4c) on the percentage of spontaneous alternation. In contrast to results at 8 months, Tukey’s multiple comparisons indicated no difference between MIC- and CTRL-fed 3xTgAD mice (p = 0.9298; Fig. 5c, Sup. Table 4c).

Consistent with the 8-month analysis, the locomotor activity assessment at 15 months revealed no significant interaction (F_(1, 50)_ = 0.801, p = 0.375, partial η² = 0.016) and no main effect of diet (F_(1, 50)_ = 0.080, p = 0.778, partial η² = 0.002), but there was a again highly significant main effect of genotype indicating reduced locomotion in 3xTg AD mice (F_(1, 50)_ = 46.216, p < 0.001, partial η² = 0.480; Sup. Table 4d). Post hoc multiple comparisons again mirrored earlier findings, showing no significant differences between MIC- and CTRL-fed mice within either genotype (3xTgAD: p = 0.9341; nonTg: p = 0.8722; Fig. 5d; Sup. Table 4d).

The reduced motor activity observed in the Y-maze was corroborated by open-field testing which measured total distance for the 15-minute duration of the test. At 8 months of age, there were no significant interaction effects (F_(1, 55)_ = 0.414, p = 0.522, partial η² = 0.007) or main effects of diet (F_(1, 55)_ = 0.002, p = 0.962, partial η² < 0.001; Sup. Table 5a). Tukey’s post hoc comparisons revealed no significant differences between diets within either genotype (nonTg, p = 0.9753; 3xTg AD, p = 0.9602; Sup. Fig. 6a, b; Sup. Table 5a). However, a significant main effect of genotype was detected, indicating reduced locomotion in 3xTg AD mice (F_(1, 55)_ = 11.615, p = 0.001, partial η² = 0.174; Sup. Table 5a). Similarly, at 15 months of age, there were no significant interaction effects (F_(1, 51)_ = 0.137, p = 0.713, partial η² = 0.003) or main effects of diet (F_(1, 51)_ = 1.844, p = 0.180, partial η² = 0.035), nor differences between diets within either genotype (nonTg, p = 0.6229; 3xTg AD, p = 0.8947; Sup. Fig. 6c, d; Sup. Table 5b). In contrast, a highly significant main effect of genotype was again observed, reflecting markedly reduced motor activity in 3xTg AD mice (F_(1, 51)_ = 43.264, p < 0.001, partial η² = 0.459, Sup. Table 5b).

**Figure 6:**
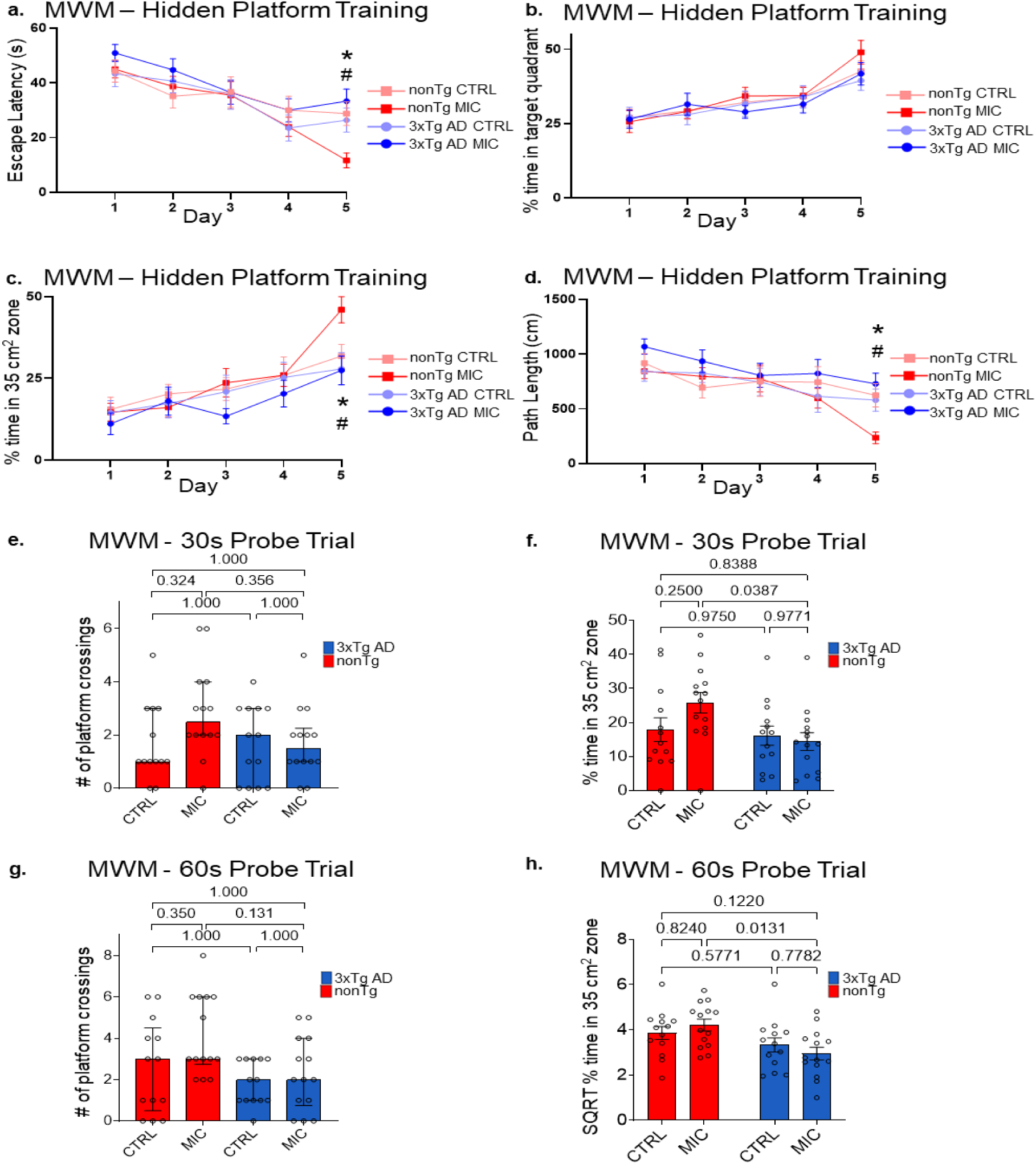
MIC improves spatial memory performance in 15-month-old nonTg mice but not 3xTg AD mice. **(a-d)** Assessment of spatial acquisition during the 5 training days for **(a)** scape latency, **(b)** percent time in target quadrant, **(c)** percent time in 35 cm^2^ target zone, and **(d)** path length. **\*** nonTg CTRL vs. nonTg MIC, **#** nonTg MIC vs 3xTg MIC, p value < 0.05. Tukey multiple comparisons: **(a)** Day 2 – Day 5**; (b)** Day 1 – 5**; (c)** Day 2 & 5**; (d)** Day 2 – 5. Bars display mean ± SEM. Bonferroni-corrected multiple comparisons on ranks: **(a)** Day 1 & 5 **(c)** Day 1,3 & 4 **(d)** Day 1. Bars display median ± interquartile range (IQR). **(e, f)** Assessment of 30 second probe trials for **(e)** number (#) of platform crossings and **(f)** percent time in 35 cm^2^ target zone. **(g, h)** Assessment of 60 second probe trials for **(g)** number (#) of platform crossings and **(h)** percent time in 35 cm^2^ target zone. **(e, g)** Tukey multiple comparisons. **(f, h)** Bonferroni-corrected multiple comparisons on ranks. If data violated assumptions of normality or homogeneity of variance, a transformation was applied and marked with a single asterisk. If transformation failed to correct these violations, non-parametric Kruskal-Wallis tests with Bonferroni-corrected pairwise comparisons were used and marked with a double asterisk For full statistical analysis, see Sup. Table 6. Significance, p < 0.05. Bars display mean ± SEM.

The combined results from the Y-maze and open-field tests demonstrate that 3xTg AD mice exhibit robust genotype-dependent reductions in exploratory and locomotor activity at both 8 and 15 months of age, consistent with the progression of neuropathology. Importantly, the MIC diet specifically improved spatial working memory performance in 8-month-old 3xTg AD mice without altering general locomotor activity, indicating a targeted cognitive benefit rather than a nonspecific enhancement of motor behavior.

### MIC enhances spatial learning and memory in nonTg mice but not 3xTg AD mice

The Morris water maze (MWM) was performed at 15 months to evaluate spatial learning and allocentric memory performance, as previously described [42, 43]. Mice were trained twice daily for five consecutive days to locate a hidden platform, followed by probe trials on day six to evaluate memory retention of the platform location in its absence.

Throughout the five-day training period, all groups progressively reduced their escape latencies, indicating effective learning (Fig. 6a). Between-group comparisons for days 1–4 revealed no significant interaction effects (p ≥ 0.162) or main effects of genotype (p ≥ 0.165) or diet (p ≥ 0.360; Fig. 6a; Supp Table 6a). On day 5, however, a non-parametric Kruskal–Wallis H test detected significant group differences (H_(3)_ = 13.757, p = 0.003, Sup. Table 6a). Bonferroni-corrected rank-based post hoc tests showed significantly shorter latencies for MIC-fed compared with control-fed nonTg mice (p = 0.025; Fig. 6a, Sup. Table 6a), indicating that MIC improved performance in nonTg but not 3xTg AD mice.

Because 3xTg AD mice displayed reduced locomotor activity in prior assays, we further evaluated performance in the MWM using measures less sensitive to motor differences. We analyzed the percentage of time spent in the target quadrant as well as within a 35 cm² zone centered on the platform location, the latter providing greater sensitivity for detecting spatial memory deficits than traditional quadrant analysis [43, 67–70]. We also evaluated the path length to the platform for training days, as well as proximity to platform for probe trials as a swimming-speed-independent index of spatial learning, minimizing potential confounds from motor impairment [43].

Percent time in the target quadrant showed no significant effects for any of the 5 training days (Fig. 6b; Sup. Table 6b). In contrast, for day 5, the more sensitive 35 cm² zone analysis showed a significant genotype effect (F_(1,50)_ = 7.011, p = 0.011, partial η² = 0.123), although without a diet effect (F_(1,50)_ = 3.604, p = 0.113, partial η² = 0.050; Fig. 6c; Sup. Table 6c). However, most of the genotype effects in day 5 seemed to be driven by better performance in nonTg MIC-fed mice (Fig. 6c, Sup. Table 6c). In fit, post-hoc tests only indicated a potentially biologically relevant increase in performance in nonTg MIC-fed mice compared to 3xTg AD MIC (p = 0.0142; Fig. 6c; Sup. Table 6c). This was further supported by a significant interaction in path-length analysis at day 5 (F_(1,50)_ = 9.867, p = 0.003, partial η² = 0.165), where MIC reduced path length in nonTg (p = 0.009) but not 3xTg AD mice (p = 0.6785; Fig. 6d; Supp Table 6d). Furthermore, the reduction in pathlength for nonTg MIC-fed mice was also significantly different from 3xTg AD MIC-fed mice (p = 0.0012; Fig. 6d; Sup. Table 6d). Taken together, these analyses indicate that group differences emerged on the fifth day of training, primarily reflecting improved performance in MIC-fed nonTg mice.

Twenty-four hours after the final training day, memory retention was assessed in 30- and 60-second probe trials by quantifying platform crossings, the percentage of time spent within the larger 35 cm² zone centered on the previous platform location, and proximity to platform. For the 30-second probe trial platform crossings, a non-parametric Kruskal-Wallis H-test did not show statistically significance differences (H_(3)_ = 5.275, p = 0.1153; Fig. 6e, Sup. Table 6e). A two-way ANOVA for assessment of the percent time in the 35 cm² target zone did not reveal a significant interaction (F_(1,50)_ = 2.611, p = 0.112, partial η² = 0.050) nor main effects for diet (F_(1,50)_ = 1.085, p = 0.303, partial η² = 0.021), but did show a significant effect for genotype (F_(1,50)_ = 4.923, p = 0.031, partial η² = 0.045; Sup. Table 6f). Albeit there was not a significant interaction, multiple comparisons indicate a potentially biologically relevant enhanced performance in MIC-fed nonTg compared to MIC-fed 3xTg AD (p = 0.0387; Fig. 6f; Sup. Table 6f), similar to the 35 cm² target zone analysis carried out for the 5^th^ training day (Fig. 6c). A two-way ANOVA assessment for proximity to platform did not reveal a significant interaction effect (F_(1,50)_ = 0.886, p = 0.351, partial η² = 0.135), but did show a significant main effect for genotype (F_(1,50)_ = 7.812, p = 0.007, partial η² = 0.135) and for diet (F_(1,50)_ = 0.037, p = 0.849, partial η² = 0.001; Sup. Table 6g). Multiple comparisons indicate trending improvement in MIC-fed nonTg compared to MIC-fed 3xTg AD (p = 0.0509; Sup. Fig. 6a; Sup. Table 6g).

For the 60-second probe trial, a Kruskal–Wallis H-test revealed significant group differences in platform crossings (H_(3)_ = 9.127, p = 0.028; Sup. Table 6h). However, Bonferroni-corrected post-hoc comparisons did not identify significant pairwise group differences (Fig. 6g; Sup. Table 6h). A two-way ANOVA examining the percentage of time spent within the 35 cm² target zone did not detect a significant interaction (F_(1,50)_ = 1.642, p = 0.206, partial η² = 0.032) or main effect of diet (F_(1,50)_ = 0.004, p = 0.951, partial η² < 0.001), but again revealed a significantly higher performance for nonTg compared to 3xTg AD (F_(1,50)_ = 9.84, p = 0.003, partial η² = 0.164; Sup. Table 6i). Although the interaction effect was not significant, as for the 30 second probe trial (Fig. 6f), post-hoc comparisons indicated a potentially biologically meaningful improvement in MIC-fed nonTg mice compared with MIC-fed 3xTg AD mice (p = 0.0131; Fig. 6h; Sup. Table 6i). A two-way ANOVA assessment for proximity to platform did not reveal a significant interaction effect (F_(1,50)_ = 1.450, p = 0.234, partial η² = 0.028), nor a significant main effect for diet (F_(1,50)_ = 0.257, p = 0.614, partial η² = 0.005) but did show a significant main effect for genotype (F_(1,50)_ = 10.648, p = 0.002, partial η² = 0.176; Sup. Table 6j). Multiple comparisons indicated significant improvement in performance in MIC-fed nonTg compared to MIC-fed 3xTg AD (p = 0.0128; Supp Fig. 6b; Supp Table 6j).

In summary, omnibus analysis of main effects for genotype in probe trials indicate better performance for nonTg over 3xTg AD mice in probe trials, as assessed by percentage of time spent in the 35 cm² target zone. However, in fit with the results found in the fifth and final day of training (Fig. 6a, c, d), multiple comparisons support a persistent genotype effect for MIC and suggested better performance in MIC-fed nonTg mice compared to MIC-fed 3xTg AD mice (Fig. 6f, g, i, j). Together, these results indicate that MIC enhances spatial learning and memory in healthy aging mice but does not reverse cognitive impairments associated with the 3xTg AD genotype at 15 months of age.

## Discussion

### Effects of MIC on Neuropathology

Analysis of pTau in hippocampal tissue revealed that MIC reduced tau pathology in the dorsal subiculum of 3xTg AD mice (Fig. 3b–3e). Leveraging 3D image reconstruction enabled reliable detection of perinuclear and intranuclear pTau species (Sup. Video 1), which would not be readily distinguishable in conventional 2D projections. Under normal conditions, nuclear tau regulates DNA stability and heterochromatin structure in both human iPSC derived neurons and mouse primary cortical neurons [55–57]. We observed intranuclear pTau in 3xTg AD mice was significantly reduce by a MIC diet (Fig. 3e). Although nuclear tau has been reported to have protective function on DNA, the mechanistic implications of MIC-mediated reductions in intranuclear pTau were not investigated here. Rather, this analysis provides a foundation for future studies examining the functional consequences of intranuclear pTau modulation in 3xTg AD mice.

Perinuclear pTau was defined as signal located within 2 µm of DAPI-labeled nuclei, capturing somatic pTau closely associated with the nuclear compartment. Thus, while the whole-field pTau intensity alone would include axonal labeling that cannot be attributed to specific cells (Fig. 3b), the DAPI-pTau proximity-based approach allowed assignment of pTau to individual cells and quantification of both the proportion of pTau-positive cells and intranuclear signal intensity. We show MIC significantly reduced both the percentage of pTau-positive cells and perinuclear pTau intensity in the dorsal subiculum (Fig. 3c, d), specifically evidencing a reduction in intracellular pTau burden in 3xTg AD mice. The early somatic accumulation of perinuclear AT8+ pTau marks the transition from axonal tau to soma-based aggregation preceding NFT formation [53, 54]. Since an MIC-supplemented diet reduced perinuclear pTau accumulation in 3xTg-AD mice, MIC may exert neuroprotective effects by reducing somatic pTau aggregation, and preventing the neurotoxicity associated with NFTs.

In a previous study we demonstrated that MIC enhances macroautophagy via upregulation of the master transcriptional regulator of lysosomal function, transcription factor EB (TFEB) [38]. In tau P301S mice, muscle-specific TFEB expression reduced both pTau immunoreactivity and Iba1+ microglial reactivity in hippocampal tissue by 9 months [71]. Similarly, viral TFEB delivery in rTg4510 tau transgenic mice reduced pTau and improved cognition through lysosomal-dependent pTau clearance [72]. Importantly, TFEB overexpression in 5xFAD mice (expressing only Aβ pathology) showed no effect on amyloid deposition, indicating TFEB’s selectivity for clearing tau over Aβ pathology. Thus, the clearance of pTau but not Aβ pathology by MIC in 3xTg AD that we have reported here aligns with evidence demonstrating TFEB-mediated clearance of Tau but not Aβ pathology.

Previous work shows 3xTg AD mice display significantly increased protein levels of microglial Iba1 compared to nonTg control animals [44, 47, 73]. In our analysis of Iba1, we found that 3xTg AD mice that received MIC had significantly reduced Iba1 total protein levels and a trending reduction in average protein levels compared to 3xTg AD CTRL mice (Fig. 4d & 4g), paralleling the pTau reduction. To determine whether this reflected direct MIC-mediated effects on microglia or secondary effects from pTau reduction, we examined aged nonTg controls. Iba1 levels remained unchanged between nonTg CTRL and nonTg MIC-treated mice, indicating MIC does not directly modulate microglial reactivity during normal aging. Considering pTau independently drives microglia reactivity in mouse models [74], it is likely that MIC-induced reductions in pTau are driving decreases in microglial activation, as evidenced in tauopathy mouse models treated with TFEB therapeutics [71, 72].

### Effects of MIC on Cognition

Consistent with previous studies reporting reduced locomotor behavior in 3xTg AD mice [75, 76], we observed that 8-month-old 3xTg AD mice displayed decreased total distance traveled in the open-field test and fewer arm entries in the Y-maze relative to nonTg controls (Supp Fig. 6a & Fig. 5b). Despite this motor impairment, MIC-fed 3xTg AD mice showed significantly higher spontaneous alternation rates compared to control-fed 3xTg AD mice, achieving nonTg control level performance (Fig. 5a). These results demonstrate that MIC improved spatial working memory at 8 months of age in 3xTg AD mice in a manner independent of locomotor activity. However, by 15 months of age the beneficial effects of MIC on 3xTgAD mice spatial working memory as assessed by the Y-maze were no longer observable (Fig. 5c). Similarly, the analysis of the MWM results at this age also showed MIC could not rescue 3xTg AD spatial learning during the 5 acquisition days (Figs. 6a - 6d) or spatial allocentric memory during the probe trials (Figs. 6e - 6h).

The age-dependent effects of MIC on 3xTg AD mice cognitive performance may be explained by our neuropathological observations. By reducing pTau and partially attenuating microglial reactivity, MIC may be able to restore cognitive function in earlier stages of diseases development. However, MIC did not reduce Aβ pathology, indicating that amyloid burden progresses unchecked. Thus, MIC may not be sufficient to offset the cumulative burden of persistent Aβ pathology and other age-related processes at later disease stages. This concept aligns with outcomes from anti-Aβ monotherapy in AD, where selective targeting of one pathology has proven insufficient to halt cognitive decline [14–16]. Our findings therefore suggest that combining MIC with Aβ-targeting therapeutics may be required to restore cognition in late-stage disease.

Even in the absence of transgenic pathology, aging itself introduces cognitive deficits [77]. Accordingly, 3xTg AD mice at 15 months likely experience both AD-driven degeneration and physiological age-related decline. Notably, while MIC did not prevent age-related cognitive loss in aged 15-month-old 3xTg AD mice, nonTg animals of the same age fed with MIC performed significantly better than nonTg controls on the final acquisition day of the Morris water maze (Fig. 6a–6d). This improvement suggests that MIC confers cognitive benefits during normal aging, independent of its earlier disease-modifying effect in 3xTg AD mice.

Together, these findings suggest that MIC exerts dual actions, enhancing spatial learning in healthy aged nonTg mice and rescuing working memory in earlier but not advanced stages of AD-like pathology in 3xTg AD mice.

### Other lysosomal enhancing compounds

Urolithin A (UA), a compound that enhances lysosomal function, upregulates lysosomal genes and improves protein degradation in SH-SY5Y cells and degrades A oligomers in mouse neuronal cells [78]. Similarly, MIC enhanced lysosomal function and improved protein degradation in mouse muscle cells [38]. When administered to 3xTg AD mice, UA reduced both Aβ and pTau [78–80], while in the present study, MIC selectively reduced pTau without affecting amyloid pathology. Studies support that Aβ pathology promotes tau pathology, but not vice versa in AD mouse models [81–84]. Thus, it is possible that UA’s primary effect was on lowering Aβ, which could have indirectly reduced pTau due to the known potentiation of tau pathology by Aβ. In contrast, MIC’s effects appear uniquely targeted toward pTau, independent of Aβ, highlighting a novel mechanistic action for treating tau pathology. These results suggest that combining agents with complementary mechanisms—such as MIC’s selective pTau targeting and UA’s ability to reduce Aβ—may provide a multifaceted and synergistic approach to AD therapy.

### Limitations and Future Directions

Here, we evaluated the effects of MIC on Aβ, pTau and microglial reactivity. Although these findings help define the specific pathological targets affected by MIC diet, additional studies will be required to characterize these effects in greater detail, and whether and how they are linked to lysosomal dysfunction. Additionally, while there are studies characterizing the function of nuclear localized Tau in *in vitro* systems [55–57], there is a lack of research on the effects of nuclear localized pTau. We propose that in future studies, researchers investigate the colocalization, or lack thereof, of pTau with nuclear protein markers to further our understanding of its effects on cellular function. We have evaluated neuropathology in aged 20-month-old mice and cognitive performance in 8 and 15-month-old mice. Evaluating both neuropathology and cognition simultaneously and at earlier timepoints may afford additional mechanistic insights into how and to which extent MIC can affect cognition.

The same mice underwent several behavioral tests. Prior studies indicate that repeated cognitive testing can improve performance [85–87], which may have masked MIC effects on cognition assessed at 15 months of age. Future experiments using behaviorally naïve mice may be necessary to determine the full cognitive impact of MIC without potential pre-training effects.

### Conclusions

Dietary supplementation of 3xTg AD mice with MIC was interrogated for its effects on neuropathology and cognition in the 3xTg AD mouse model. While MIC had little effect on Aβ, it robustly reduced pTau and had a less pronounced but significant effect on microglia reactivity. MIC also displayed dual effects on cognition, improving cognitive performance during normal aging in nonTg mice and rescuing working memory in intermediate stages of disease progression in 3xTg AD mice.

Despite FDA-approved Aβ therapies in AD, Tau pathology remains untreated and is more closely linked to cognitive decline [19]. Our results highlight MIC as a promising candidate for treating tau pathology alone or in combination with Aβ-targeting therapies, addressing an unmet need in AD.

## Acknowledgments

We thank Dr. Gordon Lithgow, Dr. Shankar Chinta, Dr. Minna Schmidt, Dr. Shika Shukla, the Andersen and Lithgow lab members as well as Stella Breslin and Harris Ingle from the Buck Institute morphology core staff for their invaluable support and contributions. We also thank Dr. Mary Sevigny from Dominican University of California. Figure 1a was prepared using BioRender. Graphs were prepared using GraphPad Prism.

## Funding

This work was funded by the National Institute on Aging of the National Institutes of Health under award number R01AG067325.

## Contributions

ZM, AR, GW, SS and AK performed the experiments. Behavior was performed by AR, GW and SS. Tissue dissection was performed by AR, GW and MC. Tissue processing and Immunohistochemistry was performed by ZM, GW and AK. Confocal imaging and image processing was performed by ZM. Behavioral analysis and statistical analysis were performed by ZM. JKA supervised the project with additional guidance from CCW and MC. CCW and JKA edited the manuscript. ZM wrote the manuscript, with input from all other authors.

## Competing Interests

The authors declare no competing interests.

## Data Availability

The datasets used and/or analyzed during this study are available from the corresponding author upon reasonable request.

## Abbreviations

3xTg AD: Triple-transgenic Alzheimer’s disease mouse model
AD: Alzheimer’s disease
ADRDs: Alzheimer’s disease and related dementias
Aβ: Amyloid-beta
pTau: Phosphorylated tau
NFTs: Neurofibrillary tangles
APP: Amyloid precursor protein
PSEN1/2: Presenilin 1/2 genes
TFEB: Transcription Factor EB
ALP: Autophagy-lysosomal pathway
NonTg: Non-transgenic (control) mice
IHC: Immunohistochemistry
MIC: Mitophagy-Inducing Compound
MWM: Morris Water Maze
Y-maze: Behavioral test for working memory
CTRL: Control diet
Iba1: Ionized calcium-binding adaptor molecule 1 (microglial marker)
DAPI: 4’,6-diamidino-2-phenylindole (nuclear stain)
PBS: Phosphate-buffered saline
TBS: Tris-buffered saline
NDS: Normal donkey serum
ROI: Region of interest
PHFs: Paired helical filaments
ASO: Antisense oligonucleotide
UA: Urolithin A
IACUC: Institutional Animal Care and Use Committee
OCT: Optimal Cutting Temperature compound
RRID: Research Resource Identifier
SEM: Standard error of the mean
ANOVA: Analysis of variance

## Notes

### Competing Interest Statement

The authors have declared no competing interest.

### Summary of Updates

Four citations were not working correctly, because they were not registered by the ctitation tool, and were not listed in the references as a result. I have corrected this. They now appear in the references as intended. We also adjusted the title from the last version to "A Compound Enhancing Lysosomal Function Reduces Tau Pathology, Microglial Reactivity, and Restores Working Memory in 3xTg AD Mice"

