## Supplemental Materials for "A Compound Enhancing Lysosomal Function Reduces Tau Pathology, Microglial Reactivity and Rescues Working Memory in 3xTg AD Mice"

#### Supplemental Figures

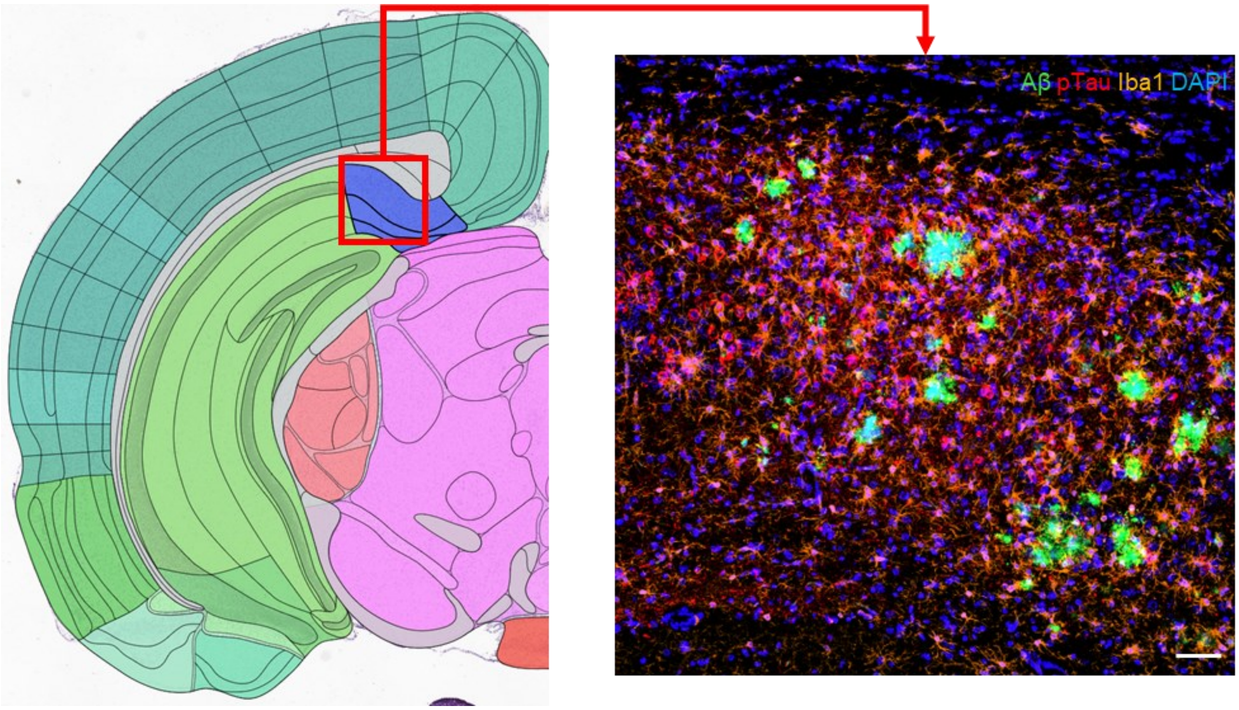


**Supplemental Figure 1: Region of interest schematic and representative stain in 3xTg AD CTRL mouse.** Region of interest schematic from Allen Brain Atlas with dorsal subiculum (blue highlight) and sampled area (red box) pointing to a representative merge image in a 3xTg AD CTRL mouse stained with Aβ (green), pTau (red), Iba1 (orange) and DAPI (blue). Scalebar = 50 μm.

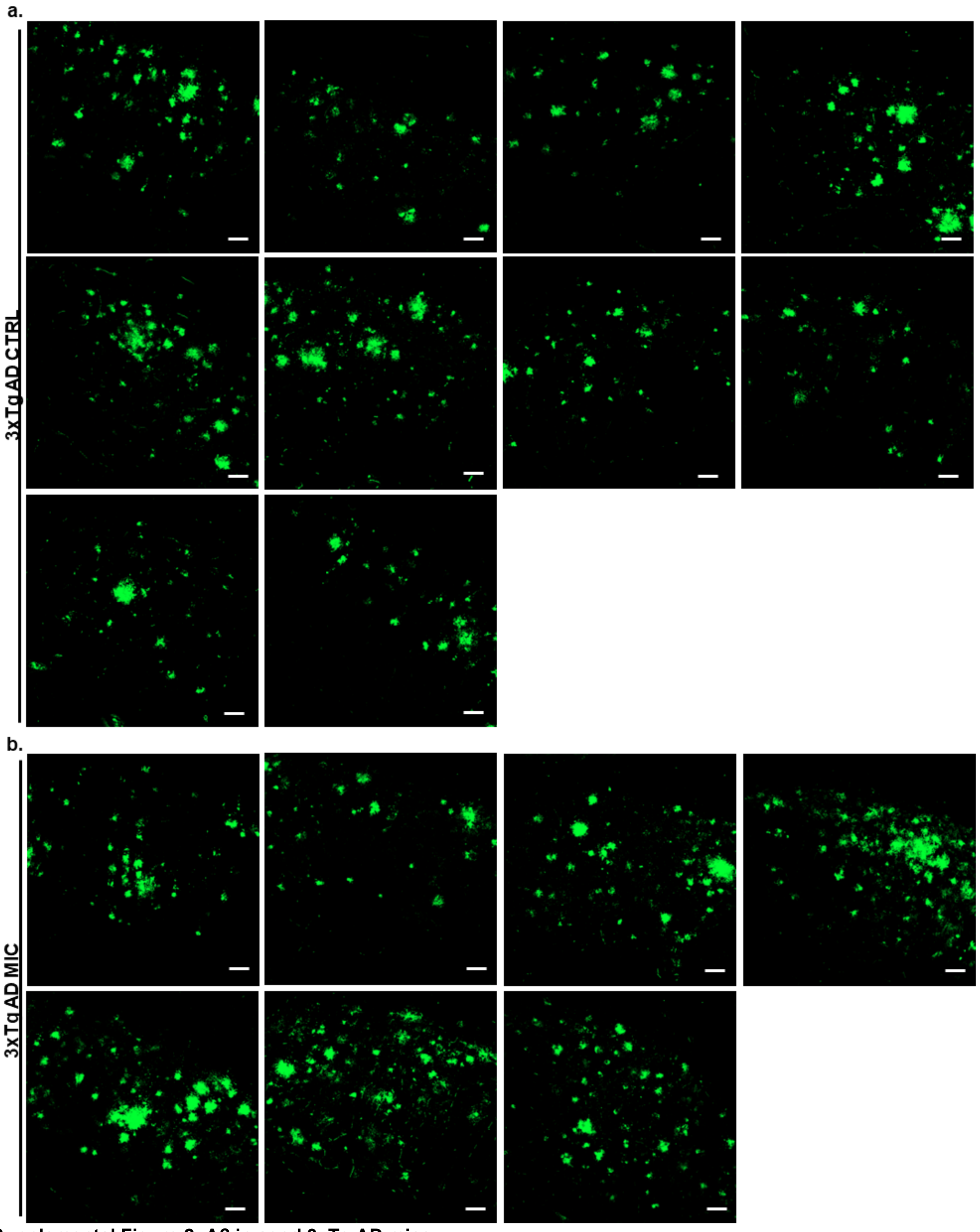


**Supplemental Figure 2: Aβ in aged 3xTg AD mice.**
Aβ (6e10, [1-16]) staining from the dorsal subiculum of 20-month-old female **(a)** 3xTg AD CTRL mice (n = 10) and **(b)** 3xTg AD MIC mice (n = 7)

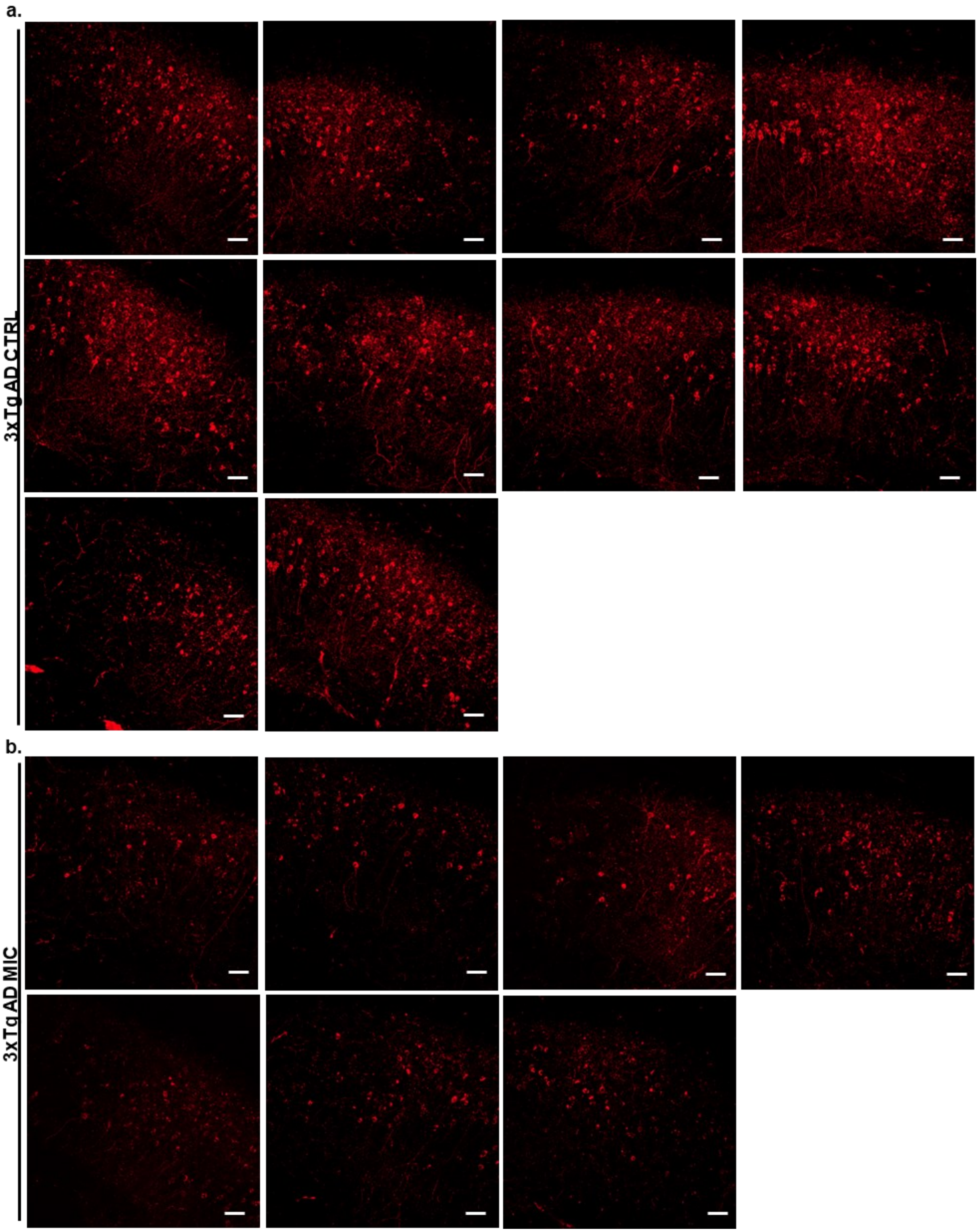


**Supplemental Figure 3: pTau in aged 3xTg AD mice**
pTau (AT8 [Serine 202/Threonine 205]) staining from the dorsal subiculum of 20-month-old female **(a)** 3xTg AD CTRL mice (n = 10) and **(b)** 3xTg AD MIC mice (n = 7)

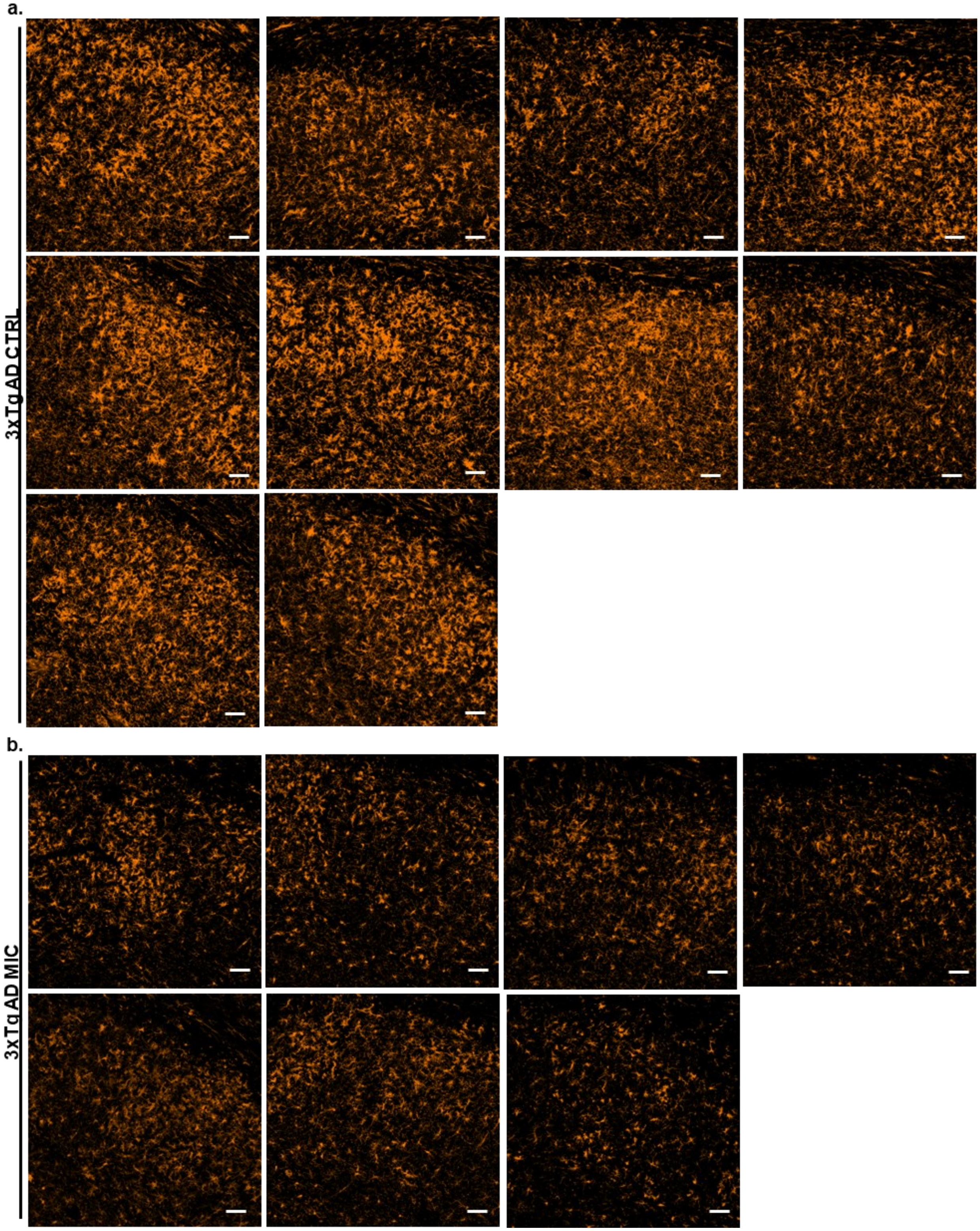


**Supplemental Figure 4: Iba1 in 3xTg AD mice**
microglia [Iba1] staining from the dorsal subiculum of 20-month-old female **(a)** 3xTg AD CTRL mice (n = 10) and **(b)** 3xTg AD MIC mice (n = 7)

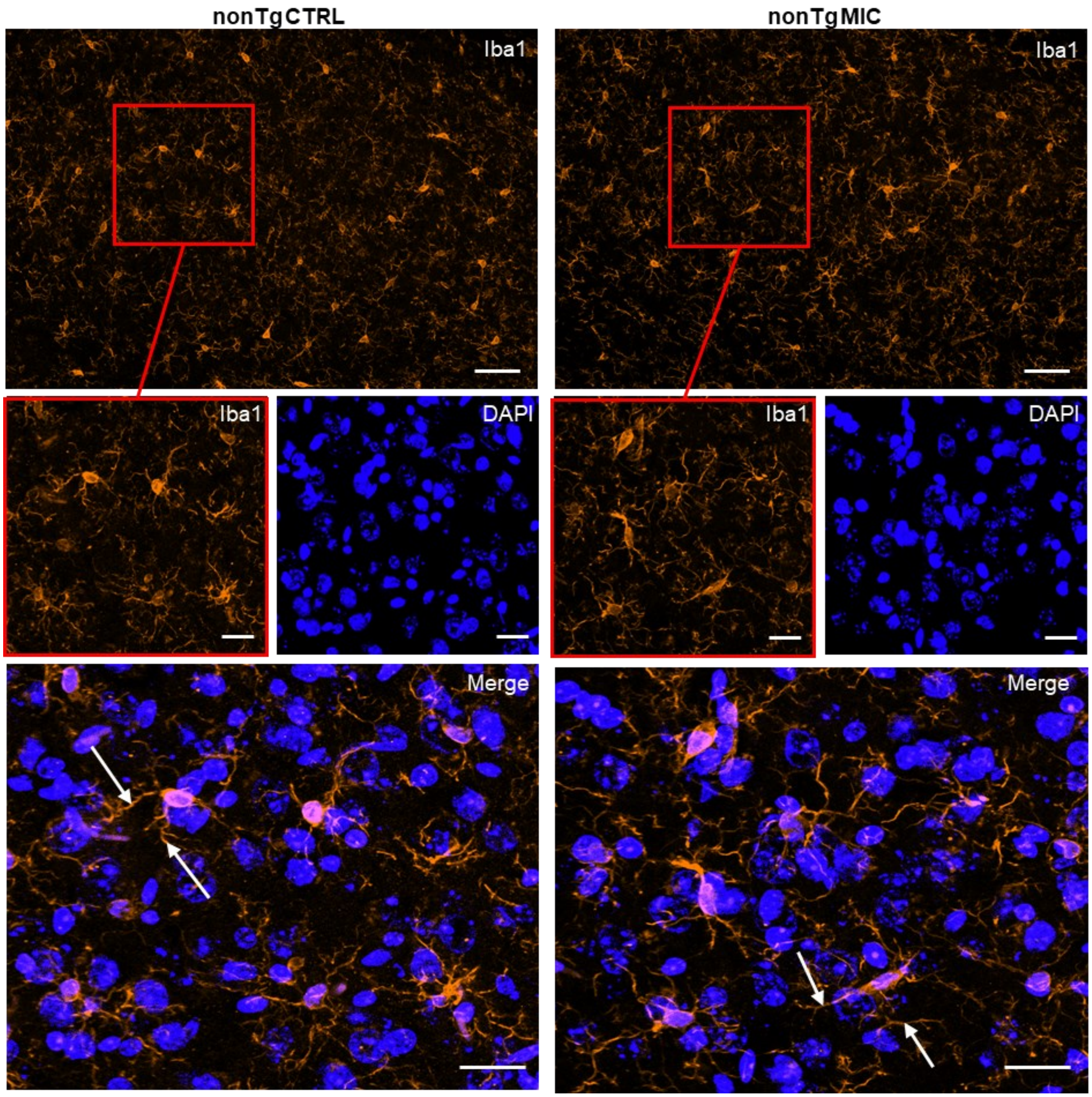


**Supplemental Figure 5: MIC treatment has no effect on microglia reactivity in dorsal subiculum of nonTg mice**

**(a)** Representative images (top) from nonTg mice on the CTRL and MIC diets visualizing Iba1 in the dorsal subiculum. Detailed images (middle) showing microglia and DAPI and merge images (bottom). Arrows to Scale bars: (top) = 50 μm; detailed images (middle) and merged images (bottom) = 20 μm.

s

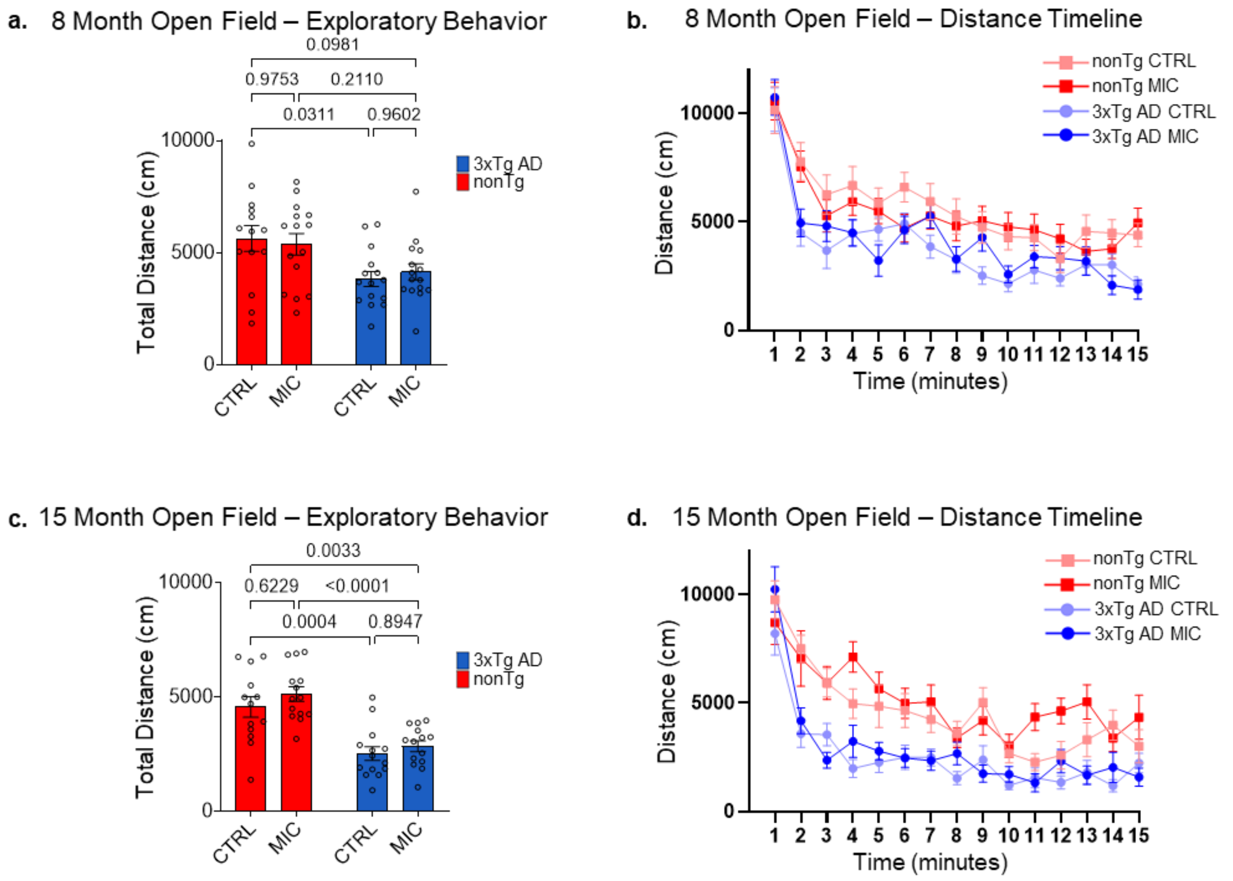


**Supplemental Figure 6: Exploratory behavior in the Open field test in 8-month and 15-month old female mice** Exploratory behavior quantified using total distance (cm) at **(a)** 8-months and **(c)** 15-months. Charts of distance traveled by minute at **(b)** 8-months and **(d)** 15-months. Values calculated from **(a, b)** 8-month old female mice, n = 14 (nonTg CTRL), n = 15 (nonTg MIC), n = 15 (3xTg AD CTRL), n = 15 (3xTg AD MIC) and **(c, d)** 15-month old female mice, n = 13 (nonTg CTRL), n = 14 (nonTg MIC), n = 14 (3xTg AD CTRL), n = 14 (3xTg AD MIC), were quantified as mean ± SEM. p-values determined using Two-way ANOVA with Tukey's multiple comparisons test, $p < 0.05$.

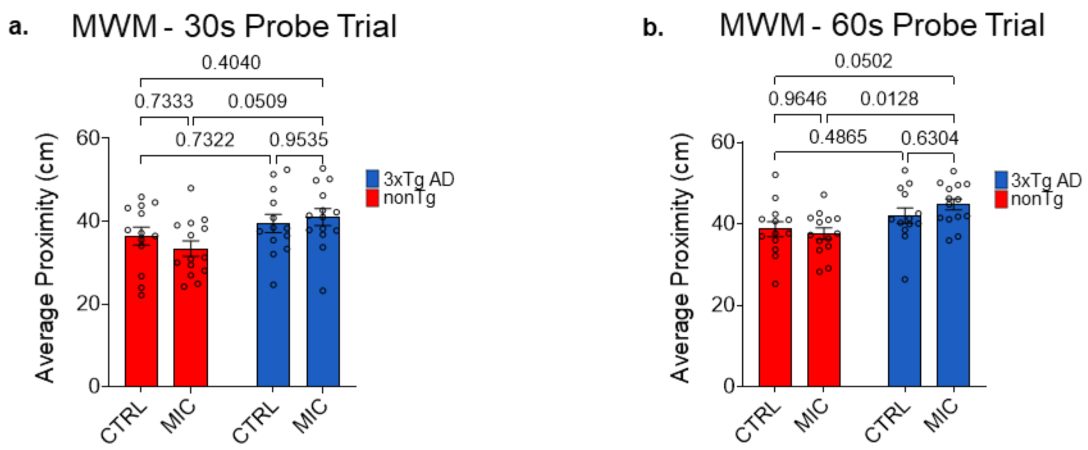


**Supplemental Figure 7: nonTg mice have better average proximity to the platform in probe trials**
Average proximity to the platform (cm) for **(a)** 30s probe trial and **(b)** 60s probe trial. All values calculated from 15-month-old female mice, n = 13 (nonTg CTRL), n = 14 (nonTg MIC), n = 13 (3xTg AD CTRL), n = 14 (3xTg AD MIC) were quantified as mean ± SEM for Two-Way ANOVA. p-values determined using Two-way ANOVA with Tukey's multiple comparisons test. Significance, $p < 0.05$.

### Supplemental Videos

**Supplemental Video 1: 3D Confocal Imaging showing pTau volume, perinuclear pTau and nuclear localized pTau at specific timestamps**

3D reconstruction from confocal Z-stack imaging of the dorsal subiculum from a 20-month-old 3xTg AD CTRL mouse with DAPI and AT8+ signal illustrating the following parameters at specific timestamps: pTau volume 00:00 – 00:09, perinuclear ptau 00:09 – 00:19, and nuclear localized pTau 00:19 – 00:29.
